# Inducible degradation of endogenous proteins by AlissAID system and development of a photoactivatable inducer

**DOI:** 10.1101/2025.02.13.638197

**Authors:** Yoshitaka Ogawa, Kohei Nishimura, Masakazu Nambo, Yuichiro Tsuchiya, Sanae Oka, Toshihiko Fujimori, Kazuya Ichihara, Akinobu Matsumoto, Keisuke Obara, Takumi Kamura

## Abstract

Protein analysis strategies involving targeted protein degradation are powerful approaches to determine gene functions. Auxin-inducible degron (AID) is among the most widely used methods for target protein knockdown. This system enables the rapid depletion of AID-tagged target proteins in an auxin-dependent manner. Various improved AID methods have been developed to date; however, the requirement to tag the target protein remains a common challenge. Here, we demonstrated the efficiency of the affinity-linker-based super-sensitive AID system for condition-knockdown of target proteins in cultured animal cells and mouse embryos. This system combines the improved AID method with small-molecule antibodies, enabling the control of GFP and mCherry fusion proteins. Additionally, this system can be used to degrade endogenous targets, such as Ras proteins. We also developed a novel inducer, caged 5-adamantyl-IAA, that precisely controlled targeted protein degradation under light irradiation. This advanced technique aids in the degradation of endogenous proteins of interest and can be used to develop new technologies for localized protein degradation.

## Introduction

Targeted protein degradation (TPD) systems are widely used for the investigating the gene function. Among of them, the auxin-inducible degron (AID) system enables to degrade the target protein in an auxin-dependent manner(*1–4*). In this system, the plant F-box protein Transport inhibitor response 1 (TIR1), which is also known as an auxin receptor, is introduced into non-plant eukaryotes. Then, TIR1 forms a SCF complex with endogenous Skp1-Cul1-Rbx1. In these organisms, AID-tag fused target protein is polyubiquitinated by SCF-TIR1 in an auxin-dependent manner and degraded by the 26S proteasome (*5, 6*). In vertebrate cell, TIR1 from *Oriza Sative* (OsTIR1) is usually used. Recently, some improved AID methods were developed to reduce cytotoxicity by excess amount of auxin (*7*). In the super-sensitive auxin-inducible degron (ssAID), OsTIR1^F74A^ mutant and 5-Adamantyl-IAA(5-Ad-IAA) is used, while an artificial pair of OsTIR1F74G mutant and another synthetic auxin, 5-Phenyl-IAA is employed. In both cases, the inducers of degradation in the nM to pM range suffice for target degradation (*8, 9*).

In the all AID method, AID-tag must be fused to target proteins for recognition. To fuse AID-tag to endogenous protein, gene manipulation is required, this is hurdled to introduce AID systems into organisms where genome editing is difficult. In addition, the basal degradation, in which AID-tagged target proteins are reduced in an auxin-independent manner, has been reported in the classical AID system (*10–13*). In some cases, target proteins fused with AID-tag are degraded even in the absence of auxin. These facts show that introducing an AID-tag is challenging when using AID system, and that the AID-tag can significantly impact the stability of the target protein. To overcome these problems, a new AID system that enables to recognize and degrade the target proteins without any degradation tags is required.

Single-domain antibodies are useful for target proteins recognising because of their small size and high specificity (*14*). Nanobody is one of the single-domain antibodies, a specialized type of antibody derived from camelid species. Unlike conventional antibodies such as IgG, Nanobodies are exceptionally small and consist of a single polypeptide chain. Some studies have reported that a fusion protein of an E3 ubiquitin ligase and nanobody induces degradation of target protein. These targeted degradation systems, such as the deGradFP system, can degrade green fluorescent proteins such as GFP recognised by nanobody: vhhGFP4, allowing existing GFP lines to be diverted to protein knockdown (*15–19*). However, the target protein degradation in these systems is triggered by the expression of the E3-nanobody fusion protein, which is difficult to strictly control.

Nanobodies have also been applied to AID systems (*20, 21*). Previously, we reported that GFP or mCherry fusion target proteins were degraded by an affinity linker-based super-sensitive AID (AlissAID) system using OsTIR1^F74A^ and AID-fused nanobodies in budding yeast (Figure 1) (*21*). In this system, 5-Ad-IAA serves as a degradation inducer, allowing for compound-induced degradation of GFP and mCherry fusion proteins at nanomolar concentrations. In addition, in the AlissAID system, basal degradation was reduced compared to that in the other AID system. However, whether the AlissAID system is effective in vertebrates remains unknown. It has also been unknown whether other small molecule antibodies than nanobodies (monobodies, designed ankyrin repeat proteins [DARPin], etc.) can be applied to the AlissAID system (*22–25*).

**Figure 1.**
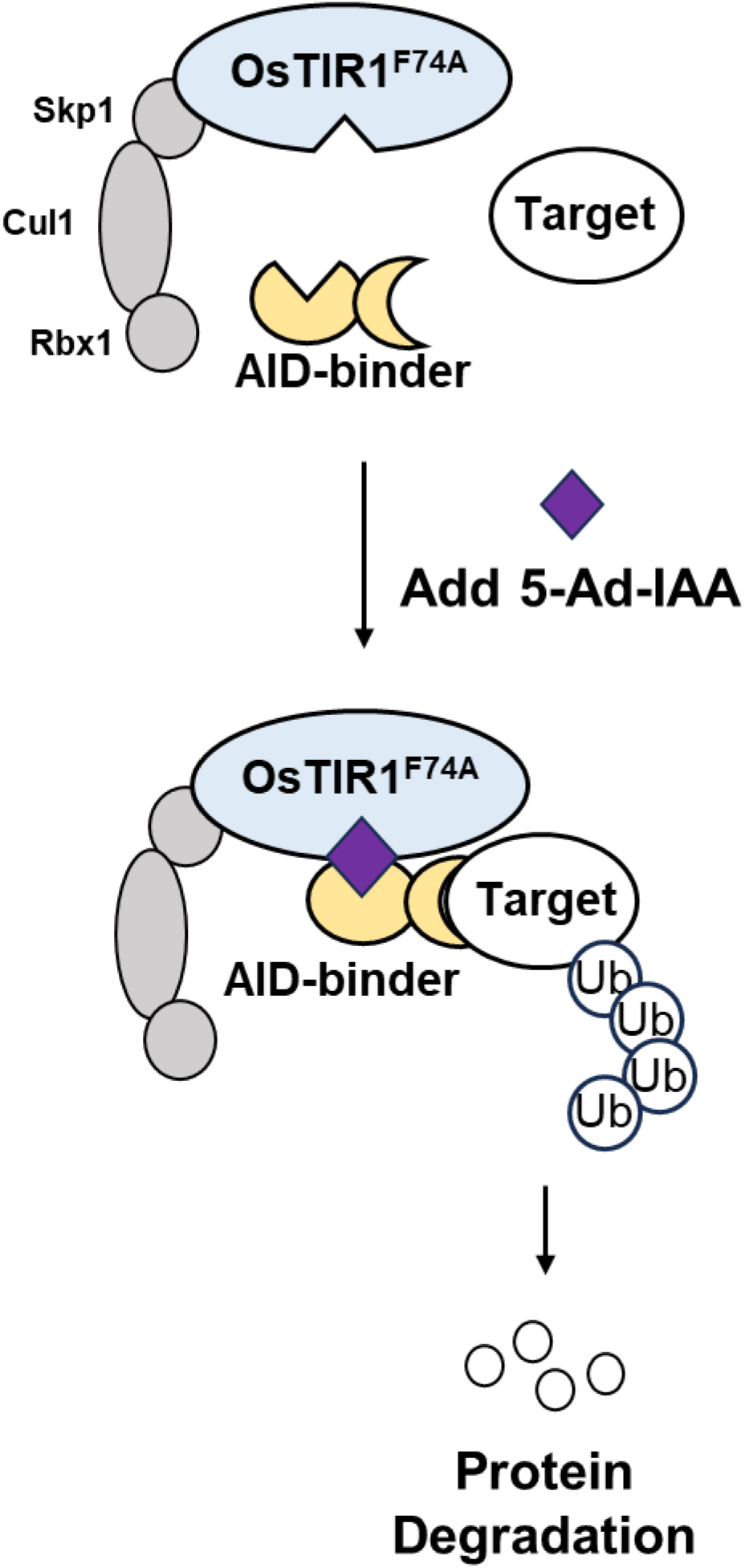
Graphical diagram of the AlissAID system. The target protein is recognized and recruited to SCF-OsTIR1^F74A^ through the AID-binder fusion protein in a 5-Adamantyl IAA-dependent manner. Subsequently, it undergoes polyubiquitination and is degraded by the 26S proteasome.

In this study, we show that the AlissAID system using nanobodies enables to degrade GFP or mCherry fused proteins effectively in various cell lines from vertebrate species, including chicken, human, and mouse. Additionally, the AlissAID system can also be used for the protein control in early mouse embryos. We show that several types of single-domain antibodies are applicable to the AlissAID system to degrade untagged endogenous proteins in the cells. Furthermore, we succeeded to develop a caged 5-adamantyl-IAA. This caged 5-adamantyl-IAA is activated by 365 nm light exposure so that the degradation of target proteins can be controlled by light exposure. The development of these advanced technologies contributes to the construction of effective protein-knockdown methods in organisms where AID-tagging is labor-intensive. Furthermore, it is expected to contribute to the development of methods for local control of target proteins.

## Result

### GFP-fused target proteins are degraded by AlissAID system in vertebrate cell lines

The AlissAID system, incorporating OsTIR1^F74A^ and vhhGFP4, degrades the target proteins fused with GFP in budding yeast (*21*). We verified whether this technique effectively functions in animal cultured cells by constructing mouse embryonic stem (mouseES) cells expressing OsTIR1^F74A^ and minimalAID-vhhGFP4^KR^ (mAID-Nb) under the EF1α promoter (Figures 1 and 2A). Consistent with previous reports, all lysine residues in mAID-vhhGFP4 were replaced with arginine residues to avoid direct ubiquitination by SCF-OsTIR1^F74A^. GFP was fused to the C-terminus of endogenous Cdk1 (cytosolic protein) or CENP-H (nuclear protein) via homology-directed repair (HDR) using clustered regularly interspaced palindromic repeat (CRISPR)/CRISPR-associated protein 9 (Cas9) genome editing. Immunoblotting revealed that both Cdk1-GFP and CENP-H-GFP rapidly degraded in a 5-Ad-IAA-dependent manner. This degradation was effectively inhibited by MG132, a proteasome inhibitor (Figure 2B; Supplementary Figure 1A). Cdk1 is involved in the progression of the M phase (*26, 27*), and CENP-H plays an important role in chromosomal segregation (*28, 29*); both of these are essential for cell proliferation. Therefore, we tested the survivability of the strains in a 5-Ad-IAA-containing medium using the colony formation assay. Proliferation of these cell lines was significantly inhibited by the addition of 5 µM of 5-Ad-IAA (Figure 2C; Supplementary Figure 1B). Furthermore, cells in which Cdk1-GFP was degraded accumulated in the G2/M phase after treatment with 5-Ad-IAA (Figure 2D). Similar experiments were performed using chicken DT40 cells, which showed 5-Ad-IAA-dependent degradation of CENP-H-GFP (Figure 2E), growth defects (Supplementary Figure 1C), and cell cycle arrest at the G2/M phase (Figure 2F). AlissAID strains expressing Cdk1-GFP or GFP-CENP-I were also generated. Endogenous Cdk1 and CENP-I were knocked out in these cells using CRISPR/Cas9. In addition, OsTIR1^F74A^, mAID-Nb, and Cdk1-GFP or GFP-CENP-I were exogenously expressed. Similar to the results of the previous experiment, we observed 5-Ad-IAA-dependent reduction in GFP-fused protein levels, decrease in GFP signals in cells, and cell cycle arrest at the G2/M phase (Supplementary Figure 2A–F). These results suggest that the AlissAID system targeting GFP is effective in various vertebrate cell lines.

**Figure 2.**
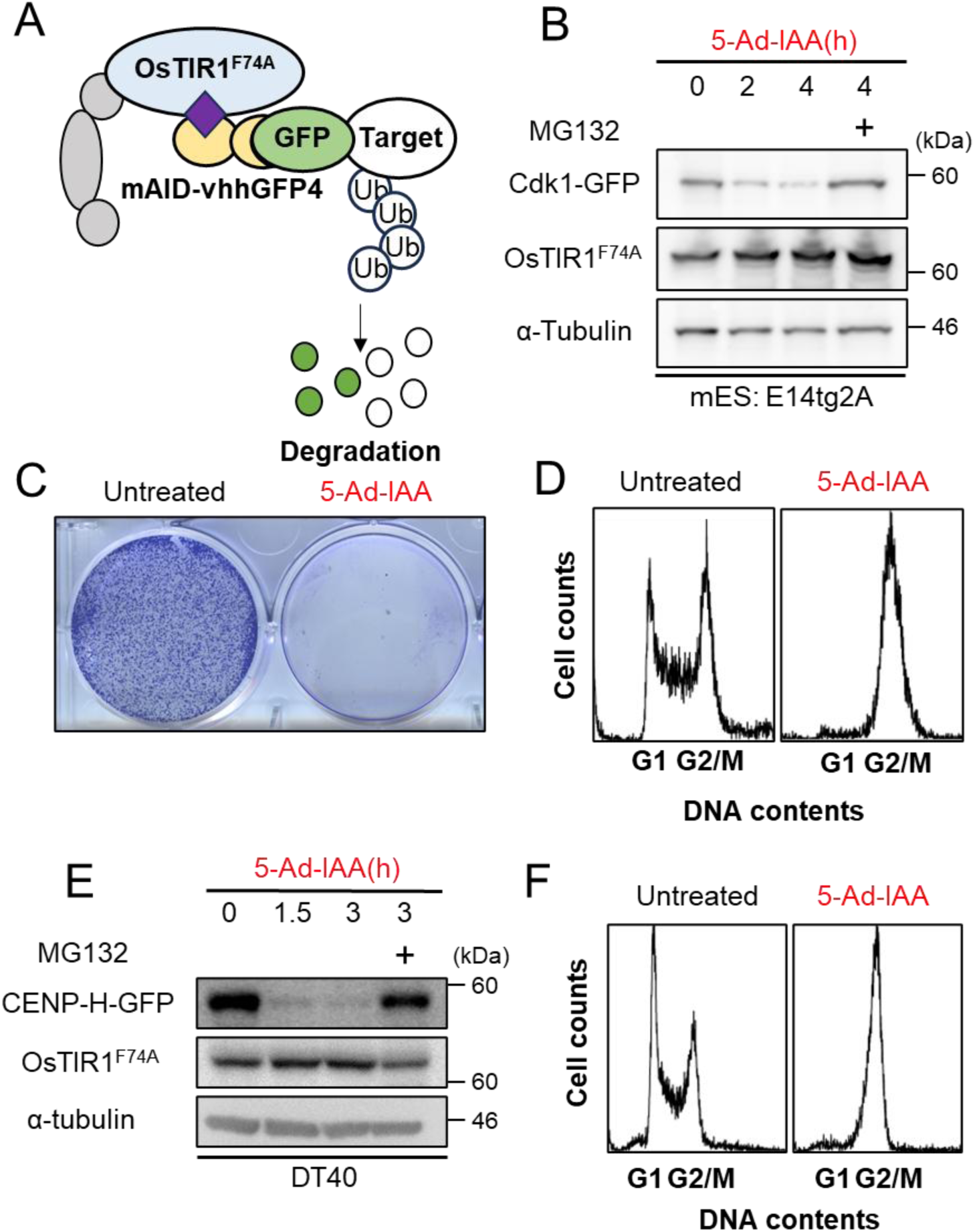
GFP fusion protein is degraded by AlissAID vhhGFP4. (A) Overview of the AlissAID system targeting GFP fusion protein. The target protein is recognized by mAID-vhhGFP4. (B) Immunoblot of *CDK1-GFP* AlissAID mouse ES cell. Cells were treated with 5 µM 5-Ad-IAA and 20 µM MG132. (C) Colony formation assay of this strain. Cells were grown in 5 µM 5-Ad-IAA containing medium and stained with crystal violet. (D) Cell cycle analysis of this strain. Cells were treated with 5 µM 5-Ad-IAA for 16 h. (E) Immunoblots of *CENP-H-GFP* AlissAID DT40 cell. Cells were treated with 5 µM 5-Ad-IAA and 20 µM MG132. (F) Cell cycle analysis of *CENP-H-GFP* AlissAID DT40 cell. Cells were treated with 5 µM 5-Ad-IAA for 16 h.

### AlissAID LaM2 effectively degrade mCherry-fused protein

Next, we tested the AlissAID system for targeting mCherry fusion proteins (Figure 3A). We constructed an AlissAID DT40 line and used KR-replaced LaM nanobodies (LaM2, LaM4, and LaM8) as binders for mCherry. mCherry was fused to the C-terminus of endogenous CENP-H via HDR using CRISPR/Cas9 genome editing. In the AlissAID system using LaM2, CENP-H-mCherry was efficiently degraded in a 5-Ad-IAA-dependent manner, and cells were arrested at the G2/M phase. In contrast, LaM4 and LaM8 did not induce the degradation of CENP-H-mCherry, and these cells did not show any cell cycle defects (Figure 3B and C). These results are consistent with those of our previous study on budding yeast (*21*).

**Figure 3.**
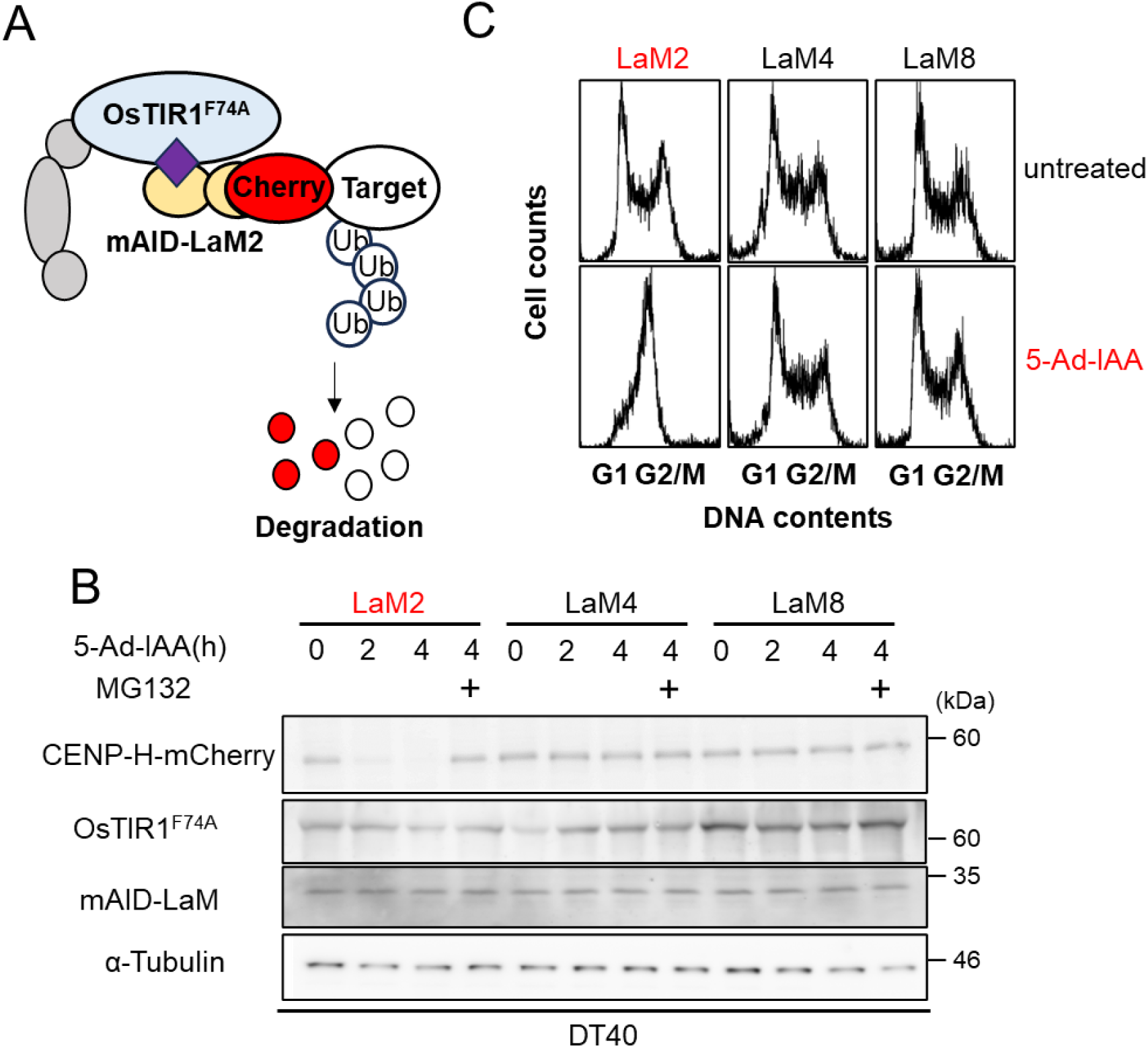
AlissAID LaM2 degrades mCherry fusion protein. (A) Overview of the AlissAID system targeting mCherry fusion protein. The target protein is recognized by mAID-LaM2. (B) Immunoblot of *CENP-H-mCherry* AlissAID DT40 cells. LaM2, LaM4 or LaM8 were used as mCherry binder. Cells were treated with 5 µM 5-Ad-IAA and 20 µM MG132. (C) Cell cycle analysis of these strains. Cells were treated with 5 µM 5-Ad-IAA for 16 h.

Various fluorescent proteins have been used in the biological field, and the specificities of the nanobodies, vhhGFP4 and LaM2, remain unclear. Therefore, we tested the specificity of the binding affinity between several fluorescent proteins (GFP, mClover3, mVenus, mScarlet, and mCherry) and nanobodies using an in vitro pull-down assay. Notably, vhhGFP4 bound to GFP, mClover3, and mVenus, but not to mScarlet and mCherry. In contrast, LaM2 bound to mCherry and mScarlet, but not to GFP, mClover3, and mVenus (Supplementary Figure 3A and B). These results suggest that these fluorescent proteins can be used as degron tags in the AlissAID systems.

### AlissAID system can induce degradation of endogenous proteins

In the AlissAID system, the target protein is recognized by a single peptide antibody, such as the nanobody; therefore, we tested the AlissAID system can target endogenous proteins by replacing the nanobody with specific antibody binding to the target protein. The Ras protein is a well-known GTPase and among thw most common oncogenic factors in humans. Here, we investigated whether the previously reported Ras binders, K55 DARPin and NS1 Monobody, (*30, 31*) can degrade endogenous Ras proteins. K55 binds to all major Ras family members: HRas, KRas, and NRas. In contrast, NS1 binds strongly to HRas and mildly to KRas, but not to NRas. First, we generated a K55-AlissAID HeLa cell line expressing both OsTIR1^F74A^ and mAID^KR^-K55 (Figure 4A). In this experiment, the KR substitution was not performed because of its potential impact on the binding affinity of K55 for Ras proteins. Immunoblot analysis showed that endogenous Ras proteins were degraded in a 5-Ad-IAA dependent manner (Figure 4B). Next, we tested the effects of NS1-AlissAID on HeLa cells. As in the case of K55, the KR substitution was not performed on NS1. Immunoblot analysis revealed that endogenous Ras proteins were degraded in a 5-Ad-IAA dependent manner (Figure 4C and D). We also generated NS1-AlissAID strains using HEK293, U2OS, and mouse embryonic stem cells. In these cells, the degradation of Ras proteins was observed (Supplementary Figure 4A, B, and C). To determine the Ras proteins targeted for degradation by the AlissAID system using NS1, we verified the binding of mAID-NS1 to HRas, KRas, and NRas using immunoprecipitation (IP) assays. Consistent with previous reports, mAID-NS1 strongly bound to HRas and weakly bound to KRas, but not NRas (Supplementary Figure 4D) (*30*). Next, we assessed the degradation of each Ras protein via flow cytometry. We generated HEK293T cells stably expressing GFP-HRas, GFP-KRas, and GFP-NRas. Additionally, we constructed the pAlissAID TT plasmid, containing the expression cassettes for OsTIR1^F74A^, mAID-binder, and mCherry. We transiently transfected the pAlissAID TT plasmid into GFP-Ras-expressing cell lines and monitored the expression levels of OsTIR1 and mAID-binder via mCherry fluorescence and Ras via GFP fluorescence. HRas was strongly degraded in a 5-Ad-IAA-dependent manner in the AlissAID system using NS1 (Figure 4E). In contrast, KRas was barely degraded, even in high mCherry-expressing cells. No degradation was observed in NRas at any level of expression. In vhhGFP4-AlissAID, used as the control, all GFP-Ras molecules were degraded. Therefore, NS1-AlissAID specifically degrades HRas, but not KRas and NRas.

**Figure 4.**
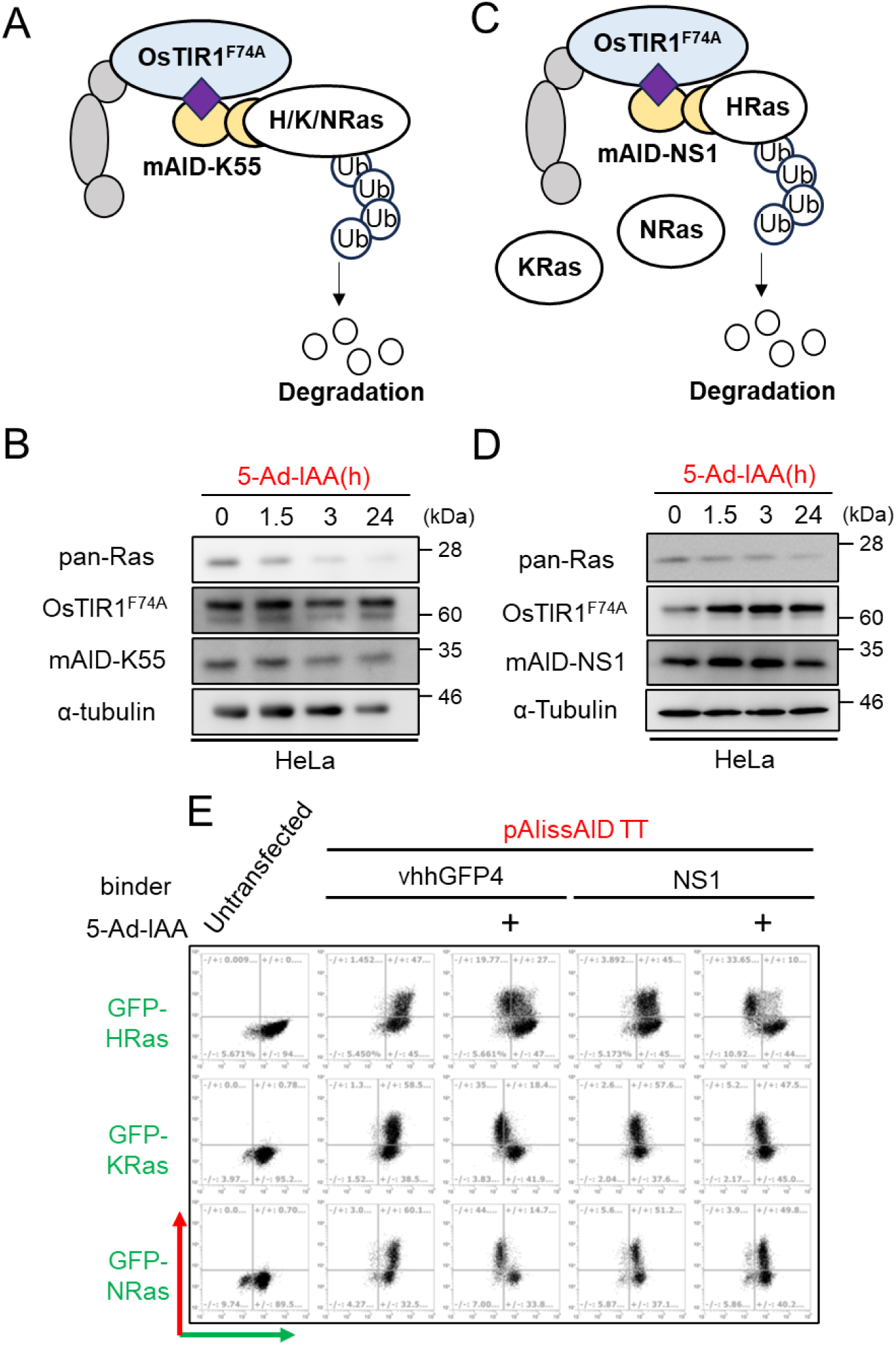
Endogenous protein degradation by AlissAID system. (A) Overview of the K55-AlissAID system. The target proteins are recognized by mAID-DARPin K55. (B) Immunoblots of K55-AlissAID HeLa cell. Cells were treated with 5 µM 5-Ad-IAA. (C) Overview of the K55-AlissAID system. The target proteins are recognized by mAID-Monobody NS1. (D) Immunoblots of NS1-AlissAID HeLa cell. Cells were treated with 5 µM 5-Ad-IAA. (E) Flow cytometry of GFP-Ras stably expression HEK293T cells transient transfected with pAlissAID TT vhhGFP4 or NS1. 5 µM 5-Ad-IAA was added to induce degradation. GFP intensity was plotted on the horizontal axis and mCherry intensity on the vertical axis, both on a logarithmic scale.

### AlissAID system facilitates phenotype analysis via endogenous protein degradation

In the previous section, we demonstrated that Ras degradation could be induced by the AlissAID system using K55, an endogenous Ras binder, and NS1. However, it is unclear whether this system induces phenotypic changes through the degradation of Ras proteins. In T24 cells harboring the HRas(G12V) mutation, proliferation decreased upon RNA knockdown of HRas (*32, 33*). Therefore, we verified HRas reduction using NS1-AlissAID in T24 cells produces a similar phenotype. First, we constructed T24 cells stably expressing OsTIR1^F74A^ and mAID-NS1 and confirmed by immunoblotting that HRas was degraded in a 5-Ad-IAA-dependent manner (Figure 5A). Next, we examined the growth of this cell line and T24 parental cells. We observed that T24 AlissAID cells exhibited significantly inhibited growth in a medium containing 5-Ad-IAA, whereas the inducer alone did not affect the proliferation of the T24 parental cell line (Figure 5B).

**Figure 5.**
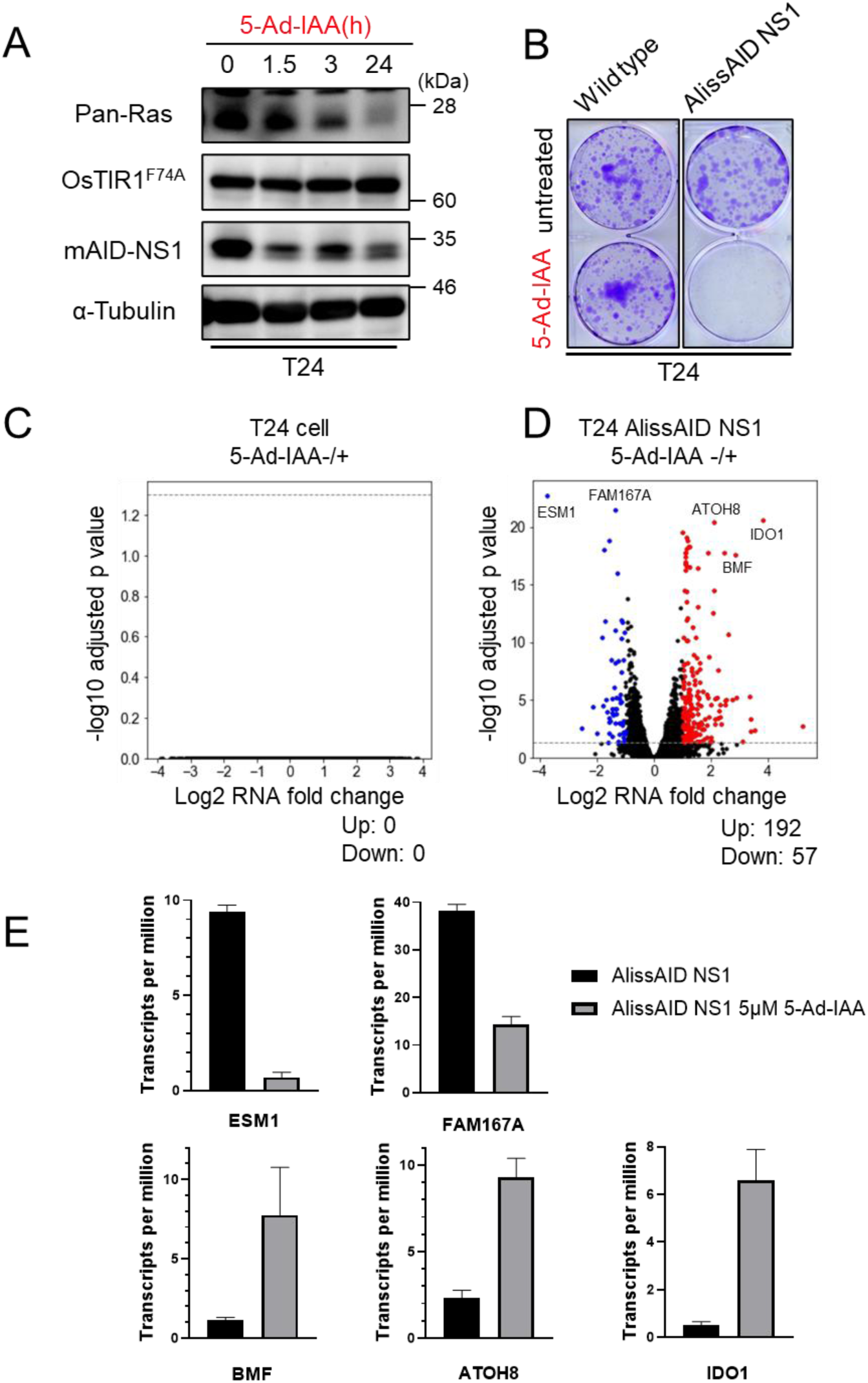
Protein degradation and phenotyping by NS1-AlissAID. (A) Immunoblot of AlissAID T24 cell. Cells were treated with 5 µM 5-Ad-IAA. (B) Colony formation assay of T24 original cell and NS1-AlissAID cell. Cells were grown in 5 µM 5-Ad-IAA containing medium, stained with crystal violet. (C, D) Volcano plot depicting log2-transformed fold change in gene expression in relation to the average of the normalized counts for each gene in the original T24 cell and T24 NS1-AlissAID cell treated with DMSO (control) or 5 µM 5-Ad-IAA for 24h (n = 3 biological replicates). (E) TPM of each gene. Error bar means ± SD.

Next, we comprehensively investigated the transcriptional responses to 5-Ad-IAA treatment and HRas degradation in cells using RNA-seq. Treatment with 5 µM 5-Ad-IAA did not affect the transcriptional status of T24 parental cells (Figure 5C). We found increased expression of some factors regulated by the Ras pathway or involved in carcinogenesis, such as IDO1, BMF, and ATOH8(*34–38*) during HRas degradation by NS1-AlissAID. In contrast, some factors showing decreased transcription levels include secretory factor ESM1, which promotes cancer cell growth, and FAM167A, which is involved in the activation of NF-kB (Figure 5D) (*39–42*). Here, gene set enrichment analysis (GSEA) revealed that the transcription of E2F target genes, which are associated with G1/S and cell cycle progression, and G2/M checkpoint genes were downregulated upon HRas degradation (Supplementary Figure 5A). These factors are involved in the Ras pathway, carcinogenesis, and cell proliferation, indicating that the degradation of HRas via NS1-AlissAID elicits the appropriate effects on cells. Therefore, degradation of endogenous proteins using the AlissAID system reduces the protein of interest (POI), thus affecting the phenotypic outcomes.

### KRas mutant-specific degradation by AlissAID system

KRAS is among the most common oncogenes mutated in various types of cancers, including non-small cell lung cancer, colorectal cancer, and pancreatic ductal adenocarcinoma. KRAS mutations are found in over 80% of all pancreatic ductal adenocarcinoma cases. G12 is a major point mutation site in KRas and key factor in cancer malignancy through changes in GTPase activity (*43, 44*). Although the development of techniques to selectively control such point mutation variants is important, recognizing point mutations using small molecules is challenging. Some single-domain antibodies exhibit low affinity for wild-type KRas but strongly bind to mutant KRas. Therefore, we investigated whether the AlissAID system, using binders targeting KRas mutants, can specifically knockdown mutant KRas. Notably, 12VC1 is a monobody of KRas(G12C) and KRas(G12V), whereas R11.1.6 is a de novo binder of KRas(G12D) mutants (*45, 46*). We transiently transfected HEK293T cells stably expressing GFP-KRas with the pAlissAID TT plasmid containing these binders and induced KRas knockdown by 5-Ad-IAA treatment. 12VC1-AlissAID degraded KRas(G12C) and KRas(G12V), but not wild-type KRas and KRas(G12D), whereas R11.1.6-AlissAID did not induce the degradation of all KRas variants, including KRas(G12D) (Figure 6A). We confirmed this result using fluorescence microscopy (Figure 6B). Overall, AlissAID system was effective for mutant variant-specific knockdown, and 12VC1 was useful for KRas G12C and G12V.

**Figure 6.**
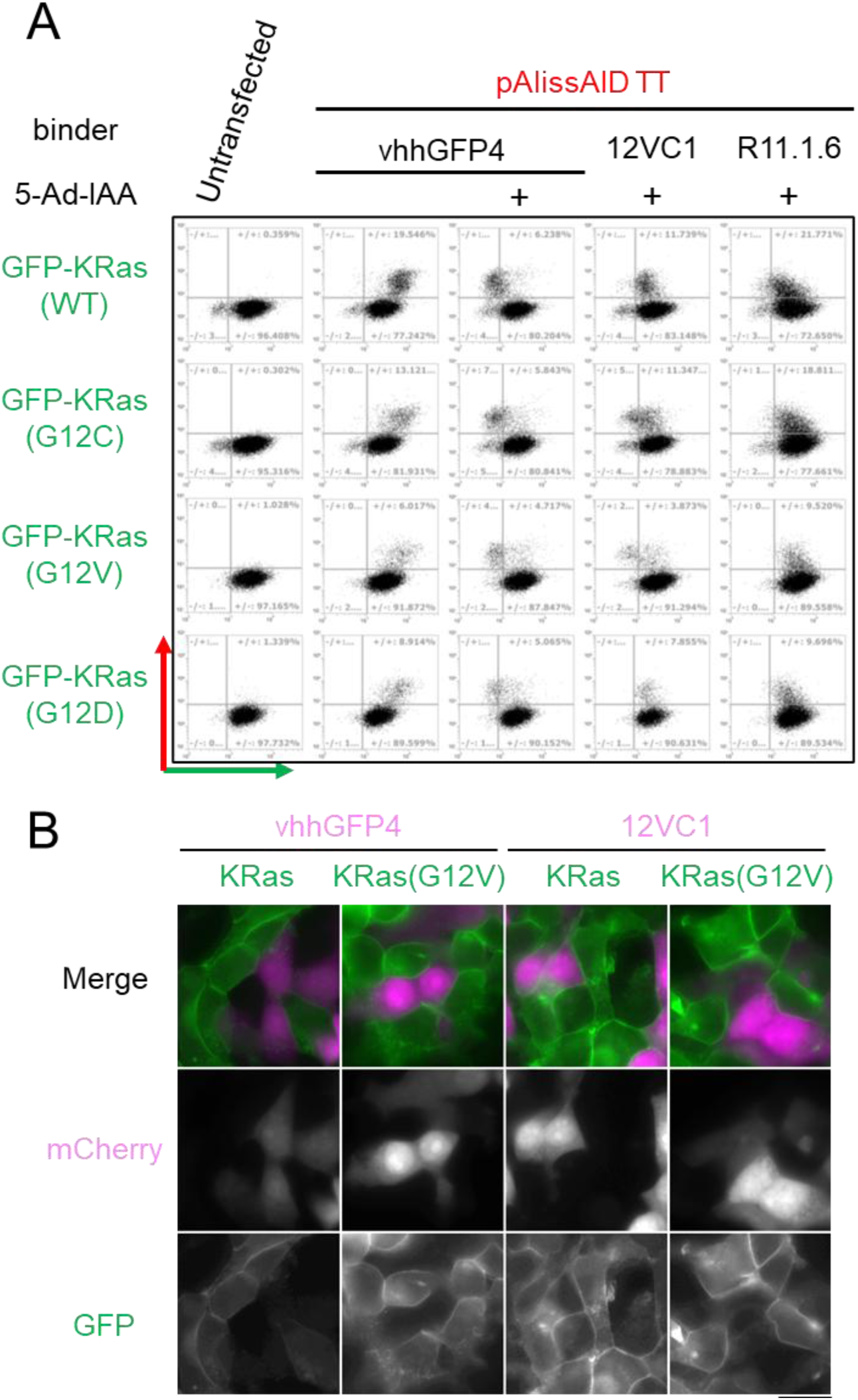
KRas mutant specific degradation by AlissAID system. (A) Flow cytometry of GFP-KRas stably expression HEK293T cells transient transfected with pAlissAID TT vhhGFP4, 12VC1 or R11.1.6. 5 µM 5-Ad-IAA was added to induce degradation. GFP intensity was plotted on the horizontal axis and mCherry intensity on the vertical axis, both on a logarithmic scale. (B) Fluorescent observation of GFP-KRas stably expression HEK293T cells transient transfected with pAlissAID TT vhhGFP4 or 12VC1. Cells were treated with 5 µM 5-Ad-IAA for 24h. Scale bar = 10 µm.

### H2B-EGFP depletion by AlissAID system in mouse embryo

Next, we evaluated the effectiveness of protein knockdown using AlissAID in early mouse embryos. Although imaging techniques using fluorescent proteins for tracking proteins are widely used, conditional protein knockdown systems, such as the AID system, are not commonly used in early mouse embryos, mainly due to the time and effort required to establish transgenic embryos with an AID-tag fused to a target protein. Therefore, protein knockdown using the AlissAID system is a useful tool for early mouse embryo analysis.

In early mouse embryos, transient expression using mRNA electroporation or microinjection has been well established. Among these, mRNA electroporation is less dependent on the skill of the experimenter and is expected to offer high reproducibility (*47*). We constructed three different plasmids for in vitro transcription, containing OsTIR1^F74A^, H2B-mCherry-P2A-mAID-Nb, or H2B-EGFP expression cassettes from pAlissAID or previously reported plasmid (*48*). Protein expression and degradation were confirmed by the transient transfection of these plasmids into HEK293T cells stably expressing GFP-KRas (Supplementary Figure 6A and B). Next, these plasmids were used as templates to synthesize mRNA via in vitro transcription. We obtained early-stage mouse embryos (E0.5) from naturally mated ICR mice and introduced mRNAs for OsTIR1^F74A^, H2B-mCherry-P2A-mAID-vhhGFP4, and H2B-EGFP by electroporation (Figure 7A). Subsequently, we used time-lapse imaging to verify whether the expression level of H2B-EGFP changed after treatment with 5-Ad-IAA. H2B-mCherry served as a non-degradable control and was used to normalize the H2B-EGFP signal. Although H2B-EGFP signals did not change in the absence of 5-Ad-IAA, H2B-EGFP signals were decreased by 5-Ad-IAA treatment (Figure 7B). This result suggests the effectiveness of conditional knockdown by the AlissAID system in early mouse embryos. The AlissAID system is thus effective as a conditional knockdown approach in organisms such as mouse embryos, where AID-tagging is labor-intensive.

**Figure 7.**
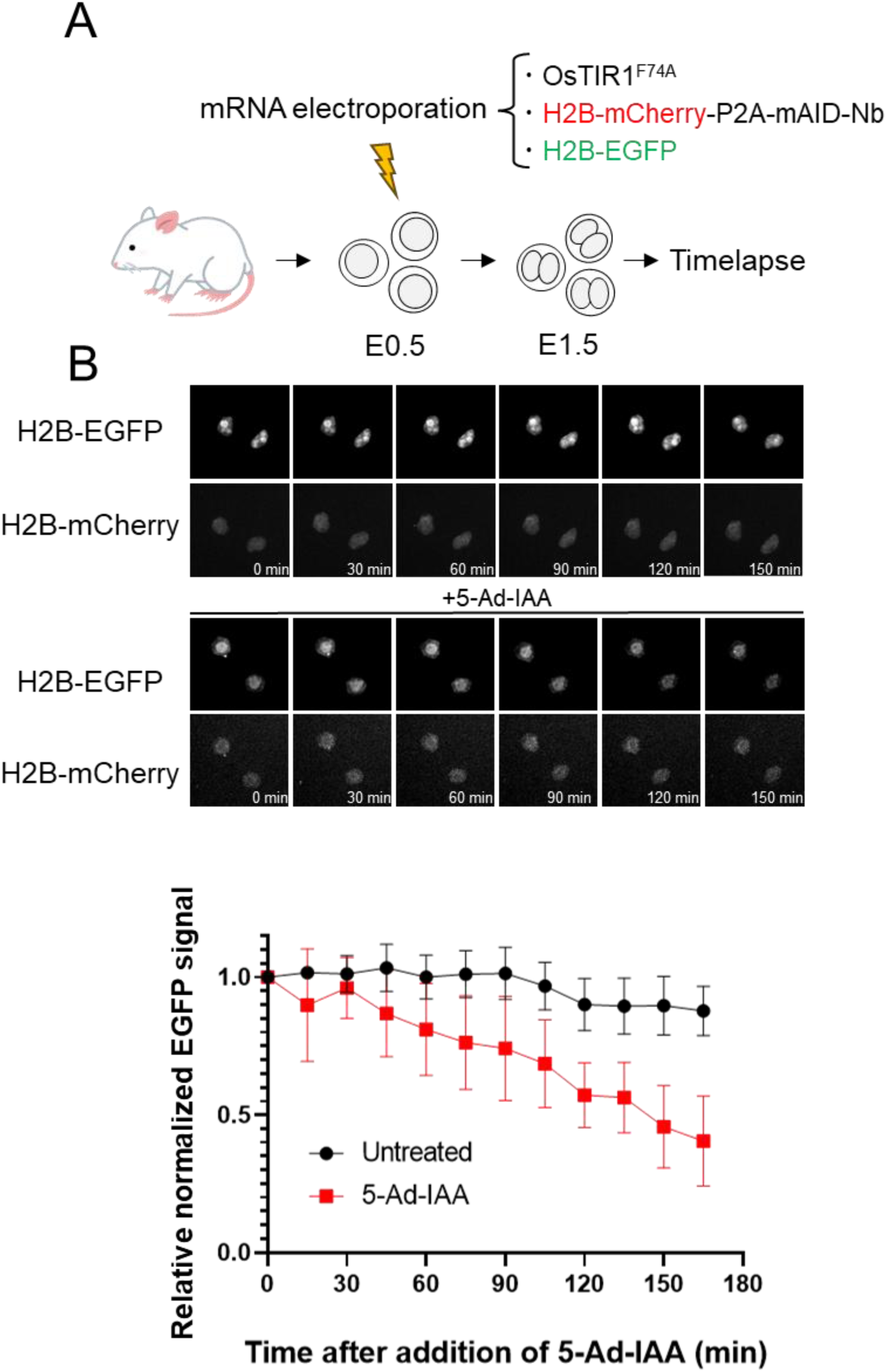
Targeted protein degradation by AlissAID system in mouse embryo. (A) Overview of the degradation assay. (B) Time lapse imaging of mouse embryo (E1.5). Embryos were treated with 5 µM 5-Ad-IAA. EGFP signals were normalized by mCherry signals and related to time 0. Erro bar means ± SD (n = 5 embryos per condition).

### Protein degradation control with photo activation of caged 5-Adamantyl IAA

Recently, opto-genetics, which controls biological phenomena using light, has been increasingly utilized in various species. Opto-genetic tools, triggered by light irradiation, allow precise control over factors such as the location, timing, and degree of activation. Attachment of photo-protection group to a functional unit of a molecule is one way to provide light-controllability. In a previous report, auxin was caged by attaching photo-protection group to the carboxylic acid moiety(*49*). We applied this method to the AlissAID system, developing an opto-genetically controlled version that enables to regulate the degradation of protein through light irradiation. In the AlissAID system, 5-Ad-IAA is used as the degrader to induce protein degradation. To achieve light-activated degradation, we synthesized a caged 5Ad-IAA (Figure 8A, Supplementary method). First, we validated the protein degradation induction activity of caged 5-Ad-IAA using a luciferase assay in ssAID system. In the absence of light exposure (Dark condition), caged 5-Ad-IAA induced minimal degradation of luciferase. However, upon activation by blue light, caged 5-Ad-IAA successfully triggered the degradation of luciferase. We compered the efficiency of luciferase degradation between 5-Ad-IAA and activated caged 5-Ad-IAA. No significant difference in the degradation induction activity was observed at concentrations ranging from 5 to 50 nM (Figure 8B). Next, we added caged 5-Ad-IAA to CENP-H-GFP AlissAID DT40 cells and observed the fluorescence of CENP-H-GFP through microscopic observation. Cells treated with caged 5-Ad-IAA showed a reduction in CENP-H-GFP fluorescence upon exposure to blue light compared with untreated cells. Conversely, no decrease in CENP-H-GFP fluorescence was observed in light non-exposed cells (Figure 8C). Finally, we validated the effectiveness of phenotype analysis using caged 5-Ad-IAA. After confirming that blue light irradiation does not affect the cell cycle (Supplementary Figure 7B), we treated the CENP-H-GFP AlissAID DT40 cells with caged 5-Ad-IAA and exposed it to blue light. As a result, accumulation in the G2/M phase of the cell cycle was observed only cells treated with caged 5-Ad-IAA and exposed to blue light (Figure 8D). This suggests that the AlissAID system utilizing caged 5-Ad-IAA is effective as an optogenetic tool for inducing light-dependent protein degradation.

**Figure 8.**
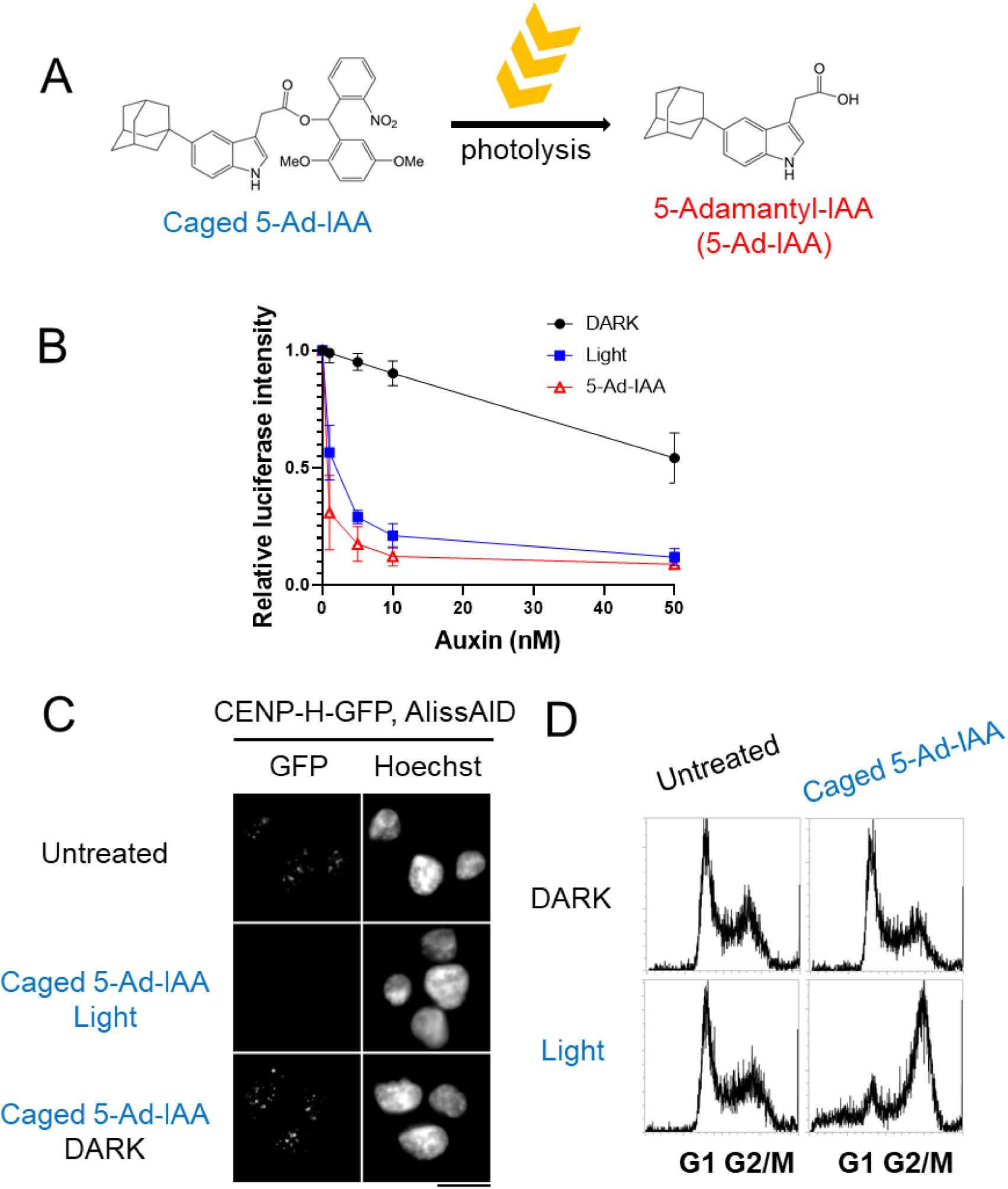
Light inducible protein degradation with caged 5-Ad-IAA. (A) Conceptual scheme depicting the photoactivation of caged 5-Ad-IAA. (B) Inducer dependent reduction of luciferase activity. Cells were treated with various concentration of auxins and 365 nm light (Red: 5-Ad-IAA, Black: caged 5-Ad-IAA in dark, Blue: caged 5-Ad-IAA and light). After 2 h treatment, luciferase activity is analyzed. Erro bar means ± SD (n = 3 biological replicates). (C) Fluorescent observation of CENP-H-GFP in AlissAID DT40 cell. Cells were treated with 50 nM caged 5-Ad-IAA and light. scale bar = 10 µm. (D) Cell cycle analysis of *CENP-H-GFP* AlissAID DT40 cell. Cells were treated with 50 nM caged 5-Ad-IAA and light.

## Discussion

IAA, a natural auxin, is widely used as an inducer to degrade AID-tagged target proteins in conventional AID systems. In contrast, super-sensitive AID and AID2 methods successfully reduce the concentration of the degradation inducer necessary for protein degradation using pairs of modified OsTIR1 and synthetic auxins. This feature was also observed in the AlissAID system, in which degradation inducer 5-Ad-IAA was effective for degradation, even at concentrations as low as 5 nM (Supplementary Figure 1C). Administration of 5-Ad-IAA at 5µM had little effect on T24 cell lines (Figure 5C), suggesting minimal side effects of 5-Ad-IAA.

Notably, a major advantage of the AlissAID system is its ability to degrade target proteins recognized by single-domain antibodies, such as nanobody. In AID systems other than the AlissAID system, AID tagging of the target protein is necessary for protein degradation, which potentially affects the function or stability of the protein. The AlissAID system addresses these issues and allows the degradation of different target proteins using various single-domain antibodies. In this study, we demonstrated that the fluorescent proteins, GFP and mCherry, can be used as degron tags for the AlissAID system. Various single-domain antibodies that recognize peptides, such as FLAG and HA, have been developed, suggesting that the AlissAID system can be adapted to use these peptides as degradation tags. This technique provides an important tool for the organisms with already established tag lines, such as a GFP-tag lines.

Another major advantage of AlissAID is its ability to degrade endogenous proteins via chemical induction. A major drawback of the AID systems is the effort required to fuse the AID tag with the target protein. Although CRISPR/Cas9 has facilitated genome editing in various eukaryotes, it is challenging to insert degradation tag sequences into all target genes through homologous recombination in polyploid cells such as cancer cells. Moreover, this process is time consuming in multicellular organisms. In the AlissAID system, the use of single-domain antibodies that directly bind to endogenous proteins can save time and effort when generating AID-based conditional knockdown lines.

Antibody molecules can bind to target proteins with high selectivity, allowing them to discern subtle differences, such as a single amino acid substitution. In this study, the 12VC1 monobody specifically degraded KRas(G12C) and KRas(G12V) (Figure 6A and B). Some antibodies recognize and bind to post-translational modifications, such as phosphorylation and acetylation. Therefore, use of such antibodies can facilitate the development of AlissAID systems specifically for the degradation of post-translationally modified proteins.

In the AlissAID system, what kind of single-domain antibody to use is crucial for the efficiency of the target protein degradation. As previously reported, AlissAID degradation assays using LaM2, LaM4 and LaM8 in budding yeast have shown that significant differences in the degradation efficiency of target proteins. Especially, the AlissAID system using LaM8 showed minimal degradation of mCherry fused proteins in both budding yeast and vertebrate cells. Similarly, R11.1.6 did not result in the degradation of the targeted KRas protein. It would be possible due to insufficient binding affinity or instability in the cell when AID-tag is fused. This remains an issue for further investigation.

Recently, various screening techniques, such as phage display, have facilitated the identification of single-domain antibodies. Therefore, a system developed to evaluate whether antibodies obtained from screening are suitable for target protein degradation will facilitate rapid selection of more effective antibodies for degradation.

Optogenetics, involving the control of molecules using light, is highly effective in animal models. For TPD, light-induced protein degradation systems, in which conformational changes of proteins are assess using the phototropic LOV2 domain, have been developed (*50, 51*). However, these systems fail to target the endogenous proteins and show basal protein reduction even without induction. Therefore, we developed a photoactivated 5-Ad-IAA degradation system in this study. Caged 5-Ad-IAA was activated by light exposure and induced target protein degradation in both the ssAID and AlissAID systems. The combination of caged 5-Ad-IAA and AlissAID system can be used for the light-induced degradation of endogenous proteins. This method is also useful for protein analysis in various organisms. Conventional AID systems are widely used for yeasts, cultured cells, flies, nematodes, and mice. Similarly, AlissAID systems could be used in various organisms. In the future, the target proteins may be freely controlled in these organisms.

## Data availability

RNA sequence data used/analyzed in this study have been deposited into the Gene Expression Omnibus database (https://www.ncbi.nlm.nih.gov/geo/) under accession number GSE267251.

## Acknowledgement

Y.O. was supported as a Research Fellow by the Japan Society for the Promotion of Science (JSPS), the Japan Science and Technology Agency (JST), or Graduate Program of Transformative Chem-Bio Research (GTR). This work was supported by JSPS KAKENHI Grant Number JP22H04926, Grant-in-Aid for Transformative Research Areas ― Platforms for Advanced Technologies and Research Resources “Advanced Bioimaging Support”. KN also received support from Kato Memorial Bioscience Foundation, Institute for Fermentation, Osaka (IFO) and Mochida Memorial foundation for Medical and Pharmaceutical Research.

## Material and methods

### Cell culture

HeLa, U2OS, HEK293, HEK293T, T24 cells are cultured in DMEM (Nakalai tesque) supplemented with 10 % FBS, 1% penicillin/streptomycin (Nekalai tesque) at 37°C in 5% CO_2_. DT40 cell are cultured in DMEM (Nakalai tesque) supplemented with 10 % FBS, 1% penicillin/streptomycin (Nekalai tesque), 1% chicken serum (Sigma), 50 μM of 2-mercaptethanol (Sigma) at 38.5°C in 5% CO_2_. Feeder free mouse embryonic stem cell (E14tg2a) are cultured on Gelatin coated dish in DMEM (Nakalai tesque) supplemented with 10% FBS, 5% Knockout Serum, 2 mM of l-glutamine (Nakalai tesque), 1% penicillin/streptomycin, 1% of non-essential amino acid (Nakalai tesque) and LIF protein at 37°C in 5% CO2. HeLa and U2OS cells were gifted from Masato Kanemaki. HEK293 and HEK293T cells were gifted from Shinya Sugiyama. T24 cells were purchased from The National BioResource Project (NBRP, https://nbrp.jp/resource/). DT40 cells were gifted from Tatsuo Fukagawa. E24tg2a cells were gifted from Tomomi Tsubouchi.

### Auxin derivative stock solutions

5-Adamantyl-IAA (Tokyo Chemical Industry: A3390) and caged 5-Adamantyl IAA (developed in this study) were diluted into dimethyl sulfoxide (DMSO) at 5 mM centration and stored at-30<C. When use, a solution of 1000 times the concentration of each final concentration was prepared and added 1/1000 to the medium.

### Plasmid construction

For gene editing, we generated the plasmid containing GFP or mCherry-T2A-Histidin D or Neomycin resistant gene between the each 1kb of homology arm in pBluescript II SL(-). For pAlissAID plasmid, codon optimized OsTIR1-T2A-Brastcidin resistant gene (BSR)-Internal Ribosomal entry site (IRES) 2-minimalized AID^KR^ tag (lysin replaced to arginine) and binders (vhhGFP4^KR^, LaM2^KR^, LaM4^KR^, LaM8^KR^, K55, NS1, 12VC1, R11.1.6) under the control of EF1a promoter are constructed in pT2/HB plasmid. pT2/HB was a gift from Perry Hackett (Addgene plasmid # 26557; http://n2t.net/addgene:26557; RRID:Addgene_26557). We used pX330-U6-Chimeric_BB-CBh-hSpCas9 (pX330) and pX335-U6-Chimeric_BB-CBh-hSpCas9n(D10A) (pX335) (Addgene #42335) for the construction of CRISPR/Cas9 vectors following reported protocol(*52*). pX330-U6-Chimeric_BB-CBh-hSpCas9 and pX335-U6-Chimeric_BB-CBh-hSpCas9n(D10A) were gifts from Feng Zhang (Addgene plasmid # 42230; http://n2t.net/addgene:42230; RRID:Addgene_42230), (Addgene plasmid # 42335; http://n2t.net/addgene:42335; RRID:Addgene_42335). For GFP tagged target gene expression, GFP fused target gene (ggCENP-I and ggCDK1) was constructed following to EF1a promoter-puro-IRES in pT2/HB plasmid. For pAlissAID TT (transient transfection) plasmid, replace of BSR to mCherry in pAlissAID plasmid. For pT2/HB plasmid genome integration, we used pCMV(CAT)T7-SB100 plasmid by co-transfection. pCMV(CAT)T7-SB100 was a gift from Zsuzsanna Izsvak (Addgene plasmid # 34879; http://n2t.net/addgene:34879; RRID:Addgene_34879) (*53*).

### Transfection and cloning

Plasmids (total 4 µg) were diluted in 50 μl of DMEM. 18 μg of Polyethylenimine (PolySciences) was diluted in 50 μl of DMEM. Each solution was mixed and incubated for 30 min at RT. The solution was add to cells (HeLa, E14tg2a) diluted in 300 µl of DMEM, vortex 5 s and incubated for 30 min at RT. After that, cells are seeded in culture dish and incubated at 37°C for 1day. For selection, medium containing 10 µg/ml blasticidin S or 1 µg/ml puromycin was used.

Plasmids (total 2 µg) were diluted in 50 μl of DMEM. 6 μg of polyethylenimine was diluted in 50 μl of DMEM. Each solution was mixed and incubated for 30 min at RT. The solution was add to cells (U2OS, HEK293, HEK293T), and incubated at 37°C for 1day. For selection, medium containing 10 µg/ml Blastcidin S or 1 µg/ml puromycin was used.

Plasmids (total 10 µg) and DT40 cells are mixed in 200 µl of DMEM, and electroporation is performed (Poring pulse: 150V, 1 msec[pulse length], 50 msec[pulse interval], 5 number of pulse, 10% decay rate +, Transfer pulse: 50V, 50 msec[pulse length], 50 msec[pulse interval], 5 number of pulse, 40% decay rate +-). After that, cells are seeded in culture dish and incubated at 38.5°C for 1day. For selection, medium containing 30 µg/ml Blastcidin S or 0.5 µg/ml puromycin or 1 mg/ml D-Histidine was used.

Plasmids (total 10 µg) and T24 cells are mixed in 100 µl of DMEM, and electroporation is performed (Poring pulse: 125V, 5 msec(pulse length), 50 msec(pulse interval), 2 number of pulse, 10% decay rate +, Transfer pulse: 20V, 50 msec(pulse length), 50 msec(pulse interval), 5 number of pulse, 40% decay rate +-). After that, cells are seeded in culture dish and incubated at 37°C for 1day. For selection, medium containing 10 µg/ml Blastcidin S was used.

### Immunoblot

Proteins were separated via SDS-PAGE and transferred to a Nitrocellulose membrane (Wako). The membrane was incubated with anti-GFP (1:2000 dilution; our laboratory), anti-RFP(1:2000 dilution; Chromo tek, 5f8-20/5f8-100), anti-TIR1 (1:2000 dilution; Medical and Biological Laboratories, PD048), anti-AID (1:2000 dilution; gifted from Prof. Karim Labib), anti-Ras (1/5000 dilution: Abcam, ab206969). HRP-conjugated anti-Rabbit IgG (1:5000 dilution; Sigma, A6154) or Peroxidase-conjugated anti-Sheep IgG (1:5000 dilution; 713-035-003, Jackson ImmnoResearch) or Peroxidase-conjugated anti-Rat IgG (1:5000 dilution; 112-035-003, Jackson ImmnoResearch) was used as the secondary anti body. Immunodetection was performed using a Luminata Forte Western HRP Substrate system (Merck Millipore, Burlington, USA, 61-0206-81) or a Chemi-Lumi One L system (Nacalai Tes que, 07880) with a bioanalyzer (LAS4000 mini; GE Healthcare Biosciences).

### Colony formation assay

Cells (200 cells) were plated to 6-well plates with or without an auxin for 1–2 weeks. Cells were fixed with cold 100% methanol for 2 min and air-dried. Cells were stained with Crystal Violette and washed with water.

### Flow cytometry

For cell cycle analysis, cells were fixed in 70% ethanol at −30°C for 1Day. Fixed cell was centrifugated, treated with 1 ml stain mix (PBS, 1 mg/ml RNase, 5 µg/ml Propidium iodide) at 37°C for 30 min. Cells were acquired by Attune Flow Cytometer (Thermo Fisher Scientific).

For degradation assay, pAlissAID TT plasmid were transfected to GFP-Ras expression HEK293T cells. After 24h, replace the medium to 5-Ad-IAA containing medium. After 24h, cells were fixed by 4% PFA in PBS and acquired by Attune Flow Cytometer.

### Cell growth assay

DT40 cells (2 × 10^4^ cells/ml) were cultured with or without an auxin derivative. Cell concentrations are calculated by Countess II (Thermo Fisher) at indicated time point after adding auxin derivatives.

### Fluorescence microscopy

Cells were grown in culture medium containing 5 µM 5-Ad-IAA or 50 nM caged 5-Ad-IAA. After centrifugation, cells were fixed by adding 4% formaldehyde in PBS for 10 min at RT. Cells were collected by centrifugation. After washing with PBS, and staining by Hoechest 33342, cells were resuspended in PBS and observed under a fluorescence microscope (AxioObserver Z1; Carl Zeiss, Oberkochen, Germany) equipped with a CCD camera (AxioCam MRm; Carl Zeiss).

### Immuno Precipitation

GFP-Ras stable expression HEK293T cells were transient transfected with the pAlissAID TT NS1 plasmid. After 48h, cells were lysed in 1ml PBS 0.2% Titon-X, centrifuged at 4℃ and the supernatant collected in a new tube. 5µl Anti-GFP GFP-Trap Dynabeads (Proteintech) were added and rotated at 4℃ for 2 hours. The beads were washed three times with cold PBS and eluted in SDS sample buffer.

### Pulldown assay

All proteins were expressed using E. coli BL21(DE3). Cells were sonicated, and then the cell lysates were incubated with GSH beads in PBS solution (GST-vhhGFP4^KR^ and GST-LaM2^KR^). Alternatively, proteins were affinity-purified using His-tag (His6-GFP, mClover3, venus, mScarlet, mCherry). The GST-vhh binding beads were dispensed, add fluorescent proteins, and rotated for 1 hour at room temperature. The beads were washed five times with PBS, and proteins were eluted by adding sample buffer.

### Luciferase assay

DT40 cells (5 × 10^5^ cells) expressing the OsTIR1^F74A^ and mScarlet-mAID-luciferase were cultured with various concentrations of inducers for 2 hours. Luciferase activity was measured with Steady-Glo Luciferase Assay System (Promega) according to a manufacture protocol.

### RNAseq

Cells were treated with 5-Ad-IAA or Mock for 24 hours, and RNA was extracted using NuclepSpin RNA plus (Takara). For library preparation, NEBNext Poly(A) mRNA Magnetic Isolation Module and NEBNext Ultra II RNA Library Prep Kit for Illumina were used, while analysis was conducted using the NextSeq 500/550 High Output Kit v2. Libraries were sequenced with Nextseq550 system (Illumina). Reads of low quality were discarded with the use of fastq_quality_trimmer and fastq_quality_filter of the FASTX-Toolkit. Ribosomal RNA reads were removed by alignment with human rRNA sequences with the use of STAR, and the remaining reads were aligned with the human transcriptome (GRCh38.p13) and human genome (hg38) also with the use of STAR. Multiple mappings were allowed. Mapped reads were counted with the use of featureCounts, and differential expression analysis was performed with the use of DESeq2. Gene set enrichment analysis (GSEA) was performed using GSEApy version 0.9.9.

### Timelapse imaging

Animal care and experiments were conducted in accordance with the Guidelines of Animal Experiment of the National Institutes of Natural Sciences. Mouse embryos were obtained from ICR female mice (SLC). The embryos were cultured in KSOM +AA media (Millipore) covered with oil at 5% CO_2_, 37°C, and subjected to microscopic imaging. Timelapse images was obtained by CV1000 system (Yokogawa) at 5% CO_2_, 37°C.

### Electroporation

OsTIR1^F74A^, H2B-mCherry-P2A-mAID-vhhGFP4, and H2B-EGFP sequences were inserted into the IVTpro template vector (Takara). These were cleaved with HIND3 and used as templates for in vitro transcription using the mMESSAGE mMACHINE™ T7 Transcription Kit (Invitrogen). The electroporation method was followed as previously reported(*47*). Briefly, each mRNA was dissolved in Opti-MEM (Gibco) at 200 ng/µl, embryos and 5 µL of mRNA solution were placed on the electrode (LF501PT1-10, BEX). Electroporation was performed (25V, 6 times, 3 ms ON +-97 ms OFF) (Genome Editor Plus, BEX) and embryos were returned to KSOM medium and incubated at 5% CO_2_, 37°C.

### Caged 5-Ad-IAA treatment and uncaging

Like 5-Ad-IAA, DMSO was used to dissolve it at a concentration of 5 mM and stored at −30°C. For use, a solution of 1000 times the final concentration was prepared and added to the medium. Uncaging was performed by 1.2x 10^5^ µJ/cm^2^ irradiating cells with a XL-1000 UV Crosslinker (UVP) equipped with BLE 8T365.

