## Supplemental data for "Inducible degradation of endogenous proteins by AlissAID system and development of a photoactivatable inducer"

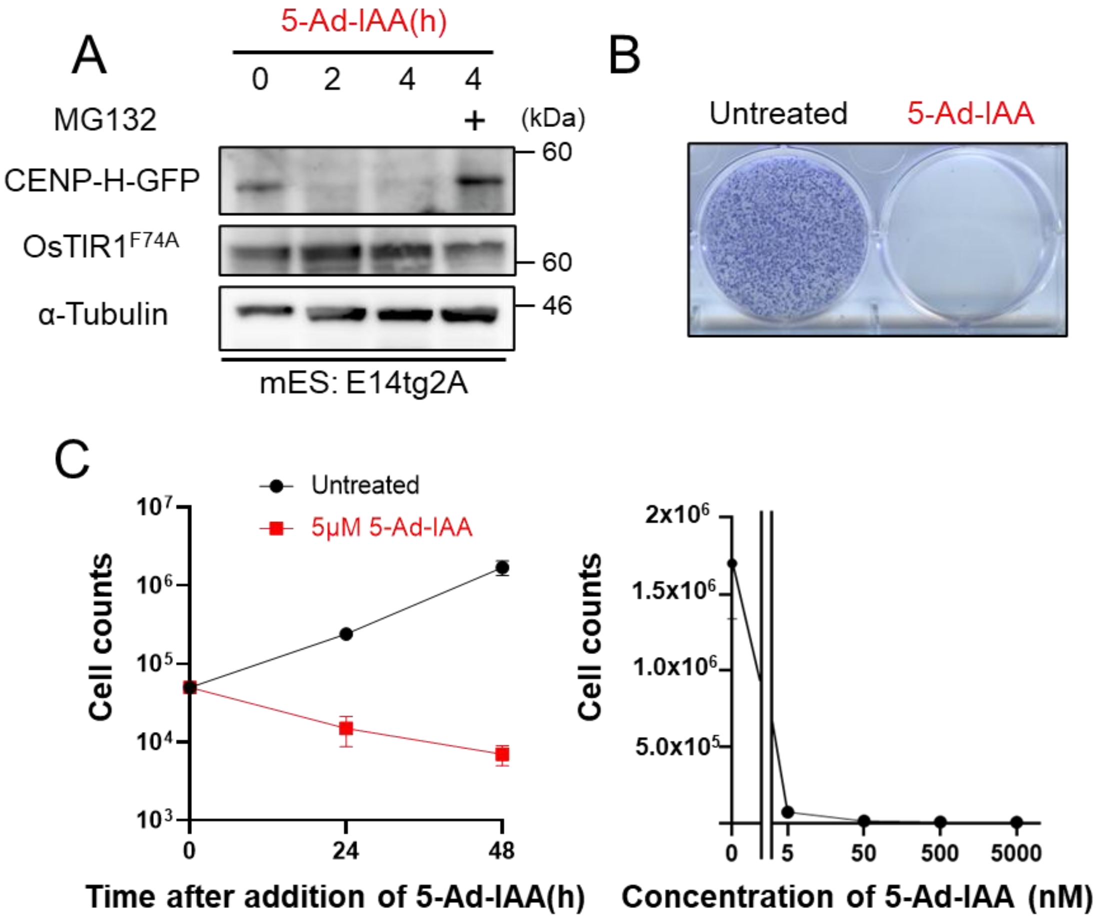


**Supplementary Figure 1. GFP fusion protein depletion by AlissAID system.**

(A) Immunoblots of *CENP-H-GFP* AlissAID mouse ES cell. Cells were treated with 5 µM 5-Ad-IAA and 20 µM MG132. (B) Colony formation assay of *CENP-H-GFP* AlissAID mouse ES cell. Cells were grown in 5 µM 5-Ad-IAA containing medium, stained with crystal violet. (C) Cell growth assay of *CENP-H-GFP* AlissAID DT40 cell. Cell counts treated with 5 µM 5-Ad-IAA at 0 h, 24 h and 48 h are shown in left graph, and cell counts with various concentration of 5-Ad-IAA at 48 h are shown in right graph. Erro bar means ± SD (n = 3 biological replicates).

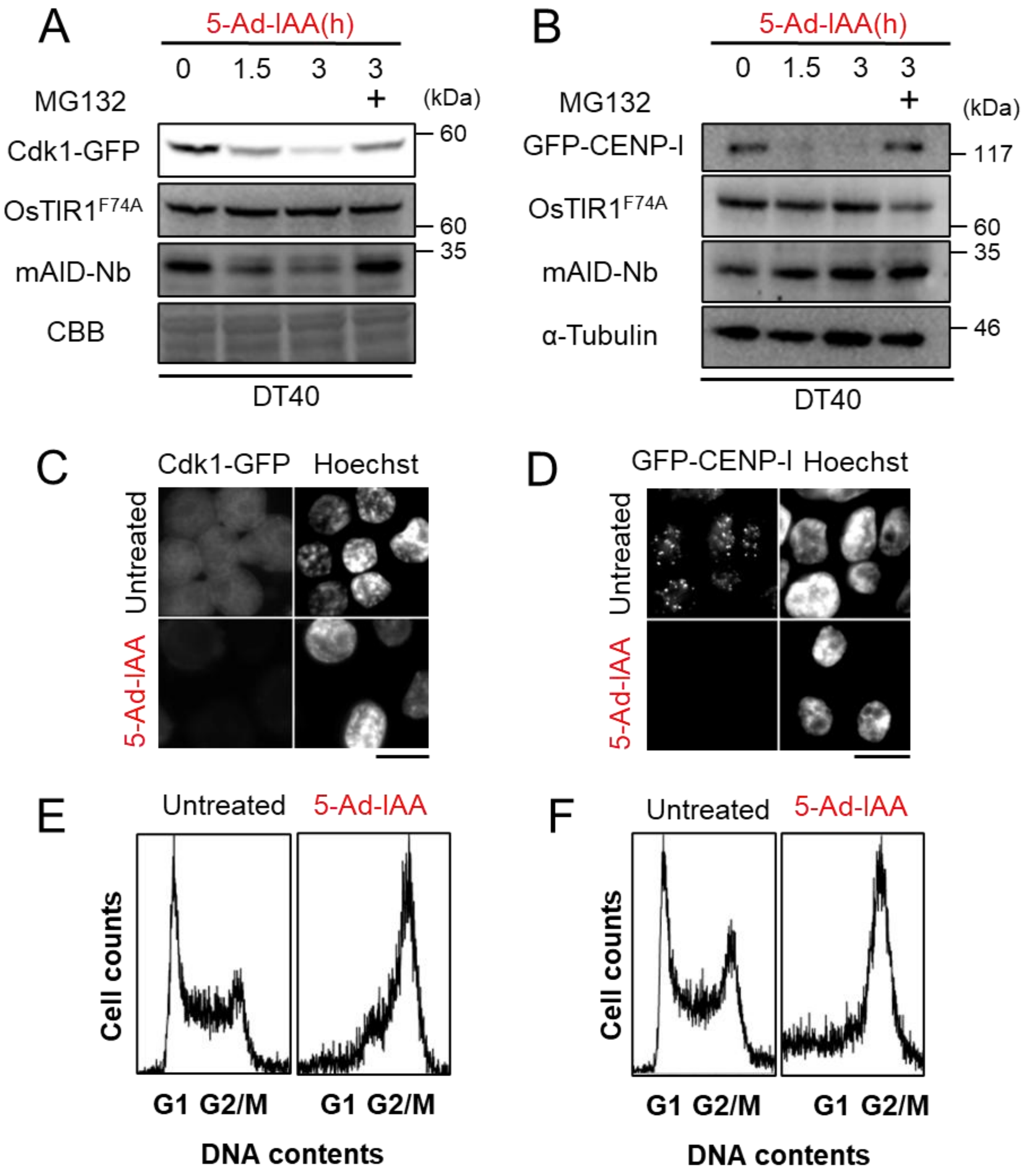


**Supplementary Figure 2. Degradation of GFP fusion protein by AlissAID vhhGFP4 in DT40 cells.**

(A, B) Immunoblots of AlissAID strains (*CDK1-GFP, GFP-CENP-I*). Cells were treated with 5 μM 5-Ad-IAA and 20 μM MG132. (C, D) Fluorescent observation of CDK1-GFP (C), GFP-CENP-I (D) in AlissAID DT40 cell. Cells were treated with 5 μM 5-Ad-IAA for 3 h. Scale bar = 10 μm (E, F) Cell cycle analysis of AlissAID strains (*CDK1-GFP* (E), *GFP-CENP-I* (F)). Cells were treated with 5 μM 5-Ad-IAA for 16 h.

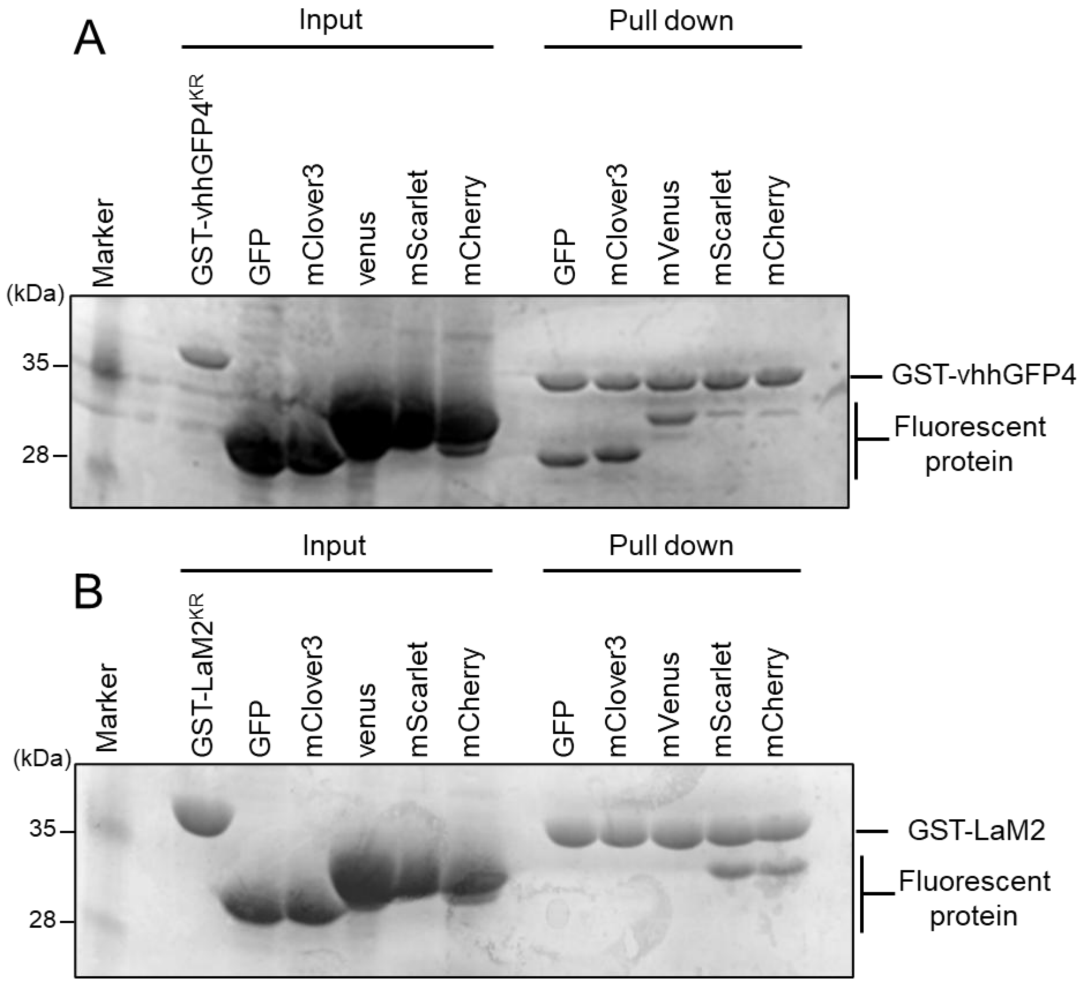


**Supplementary Figure 3. Binding potential of vhhGFP4 and vhhLaM2 to Fluorescent Proteins in vitro.**

(A, B) Pulldown assay of GST-vhhGFP4$^{KR}$ and GST-vhhLaM2$^{KR}$. The nanobodies bound to beads were incubated with His-tag-purified fluorescent proteins (GFP, mClover3, mVenus, mScarlet, mCherry), and after washing, the bound proteins on the beads were confirmed by SDS-PAGE and Coomassie Brilliant Blue staining.

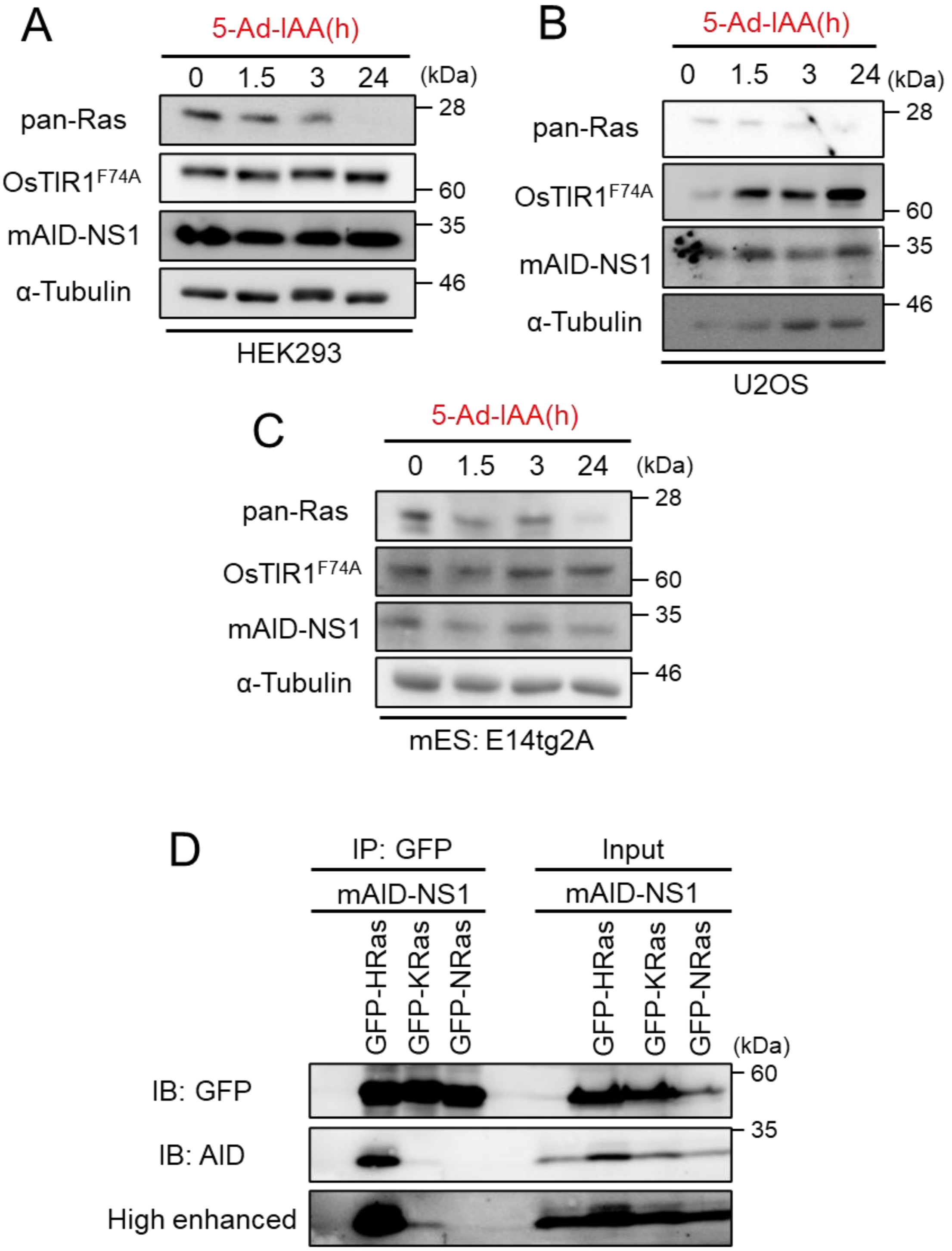


**Supplementary Figure 4. HRas specific degradation in various mammalian cells.**
(A-C) Immunoblots of HEK293 (A), U2OS (B), mouse ES (C) AlissAID NS1 cell lines. Cells were treated with 5 μM 5-Ad-IAA. (D) Immuno precipitation of GFP-Ras expression HEK293T cell. mAID-NS1 was transiently transfected for 48h.

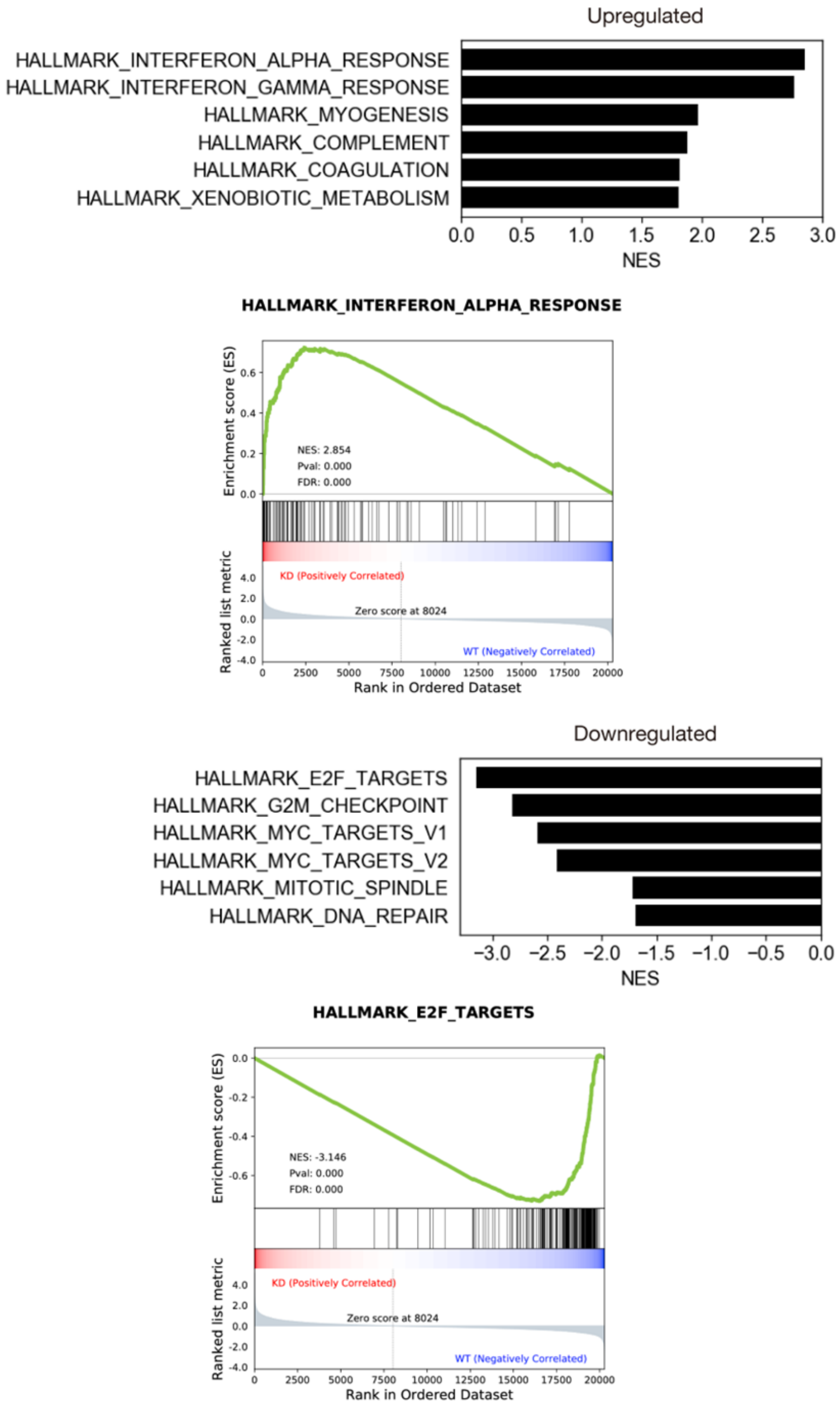


**Supplementary Figure 5. GSEA analysis of transcriptome.**

Results of pathway analysis revealing transcriptional changes identified through GSEA.

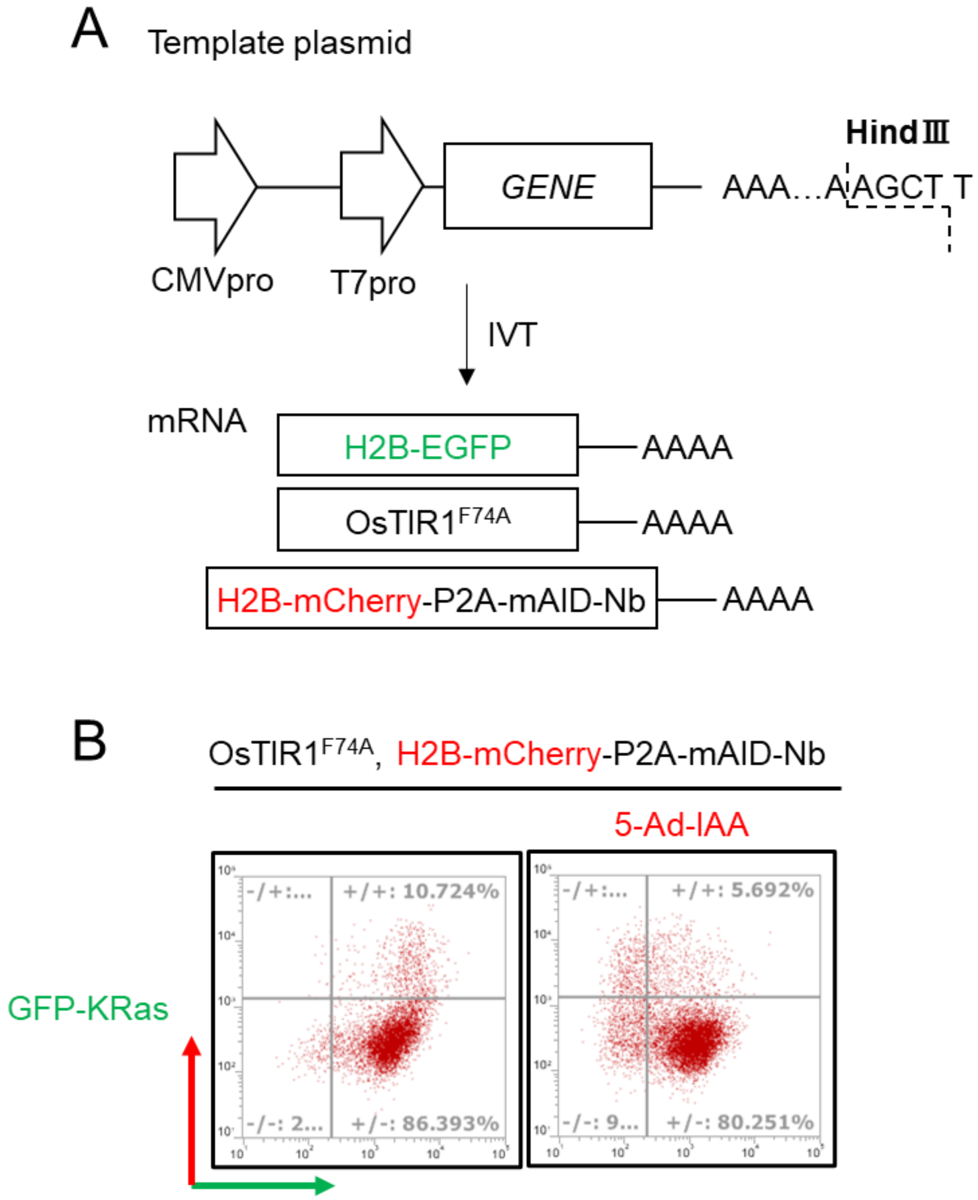


**Supplementary figure 6. IVT plasmid construction and check.**

Plasmid overview of used for IVT. (B) Flow cytometry of GFP-KRas stably expression HEK293T cell transiently transfected with IVT OsTIR1F74A and H2B-mCherry-P2A-mAID-Nb. 5 µM 5-Ad-IAA was added to induce degradation. GFP intensity was plotted on the horizontal axis and mCherry intensity on the vertical axis, both on a logarithmic scale.

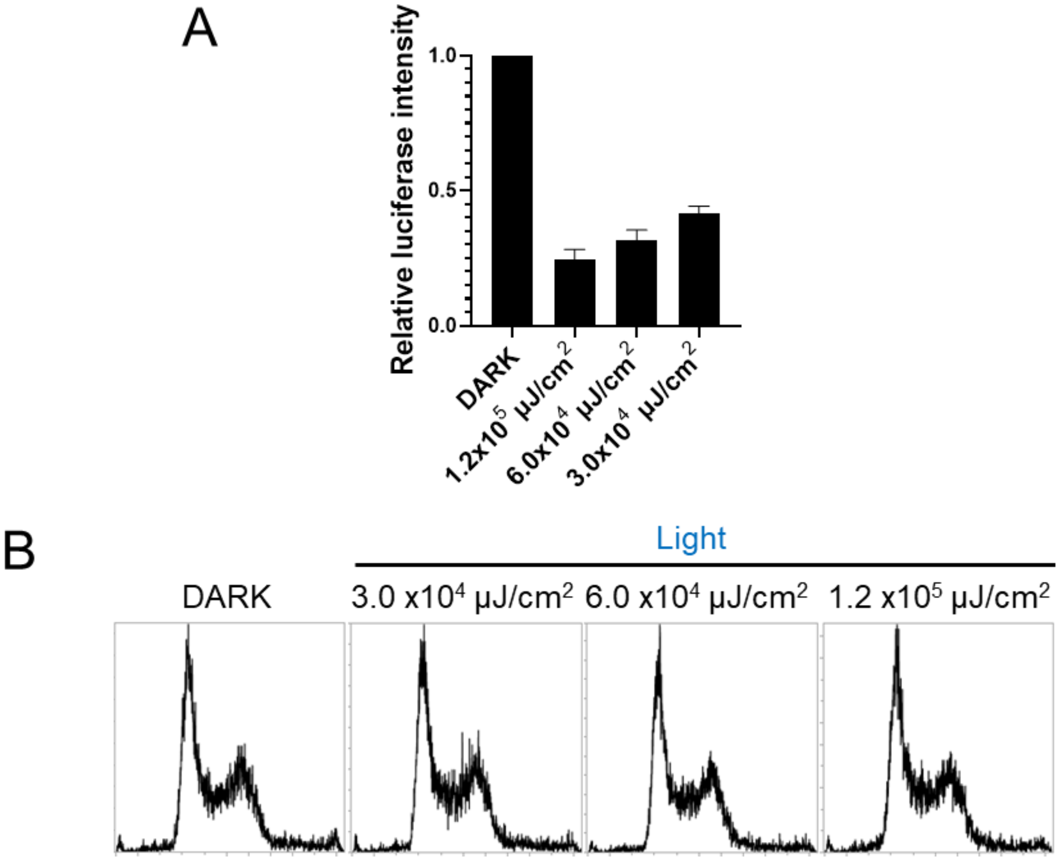


**Supplementary Figure 7. Target protein degradation with caged 5-Ad-IAA.**

(A) Luciferase activities of caged 5-Ad-IAA treated cells. The caged 5-Ad-IAA was activated using various intensities of 365 nm light. Degradation was induced for 2 hours. (B) Cell cycle analysis of DT40 cells treated 365 nm light. Analysis was conducted 24 h after light exposure.

**Supplementary method**

**Caged 5-Ad-IAA**

**1. General**

Unless otherwise noted, all materials including dry solvents were obtained from commercial suppliers and used without further purification. 2,5-dimethoxy-2'-nitrobenzhydrol was prepared according to procedures reported in the literature[49]. Unless otherwise noted, all reactions were performed with dry solvents under an atmosphere of argon in flame-dried glassware with standard vacuum-line techniques. All work-up and purification procedures were carried out with reagent-grade solvents in air.

Analytical thin-layer chromatography (TLC) was performed using Merck silica gel 60 $F_{254}$ precoated plates (0.25 mm) visualizing with UV light (254 nm) and ethanolic phosphomolybdic acid. Silica-gel for column chromatography was purchased from KANTO.

High-resolution mass spectra (HRMS) were obtained from a Thermo Fisher Scientific Exactive (ESI). Nuclear magnetic resonance (NMR) spectra were recorded on a JEOL ECA600II ($^1$H 600 MHz, $^{13}$C 150 MHz) spectrometers. Chemical shifts for $^1$H NMR are expressed in parts per million (ppm) relative to tetramethylsilane (δ 0.00 ppm). Chemical shifts for $^{13}$C NMR spectra are expressed in ppm relative to $CDCl_3$ (δ 77.00 ppm). Data are reported as follows: chemical shift, multiplicity (s = singlet, d = doublet, t = triplet, m = multiplet), coupling constant (Hz), and integration.

**2. Preparations of caged 5-Ad-IAA**

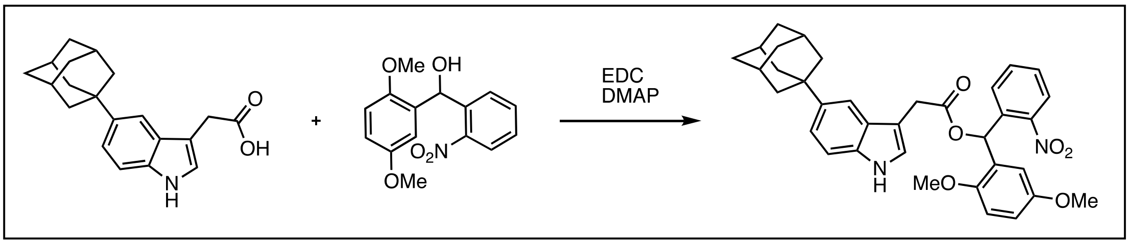


A 10 mL flask containing a magnetic stirring bar was flame-dried under vacuum and filled with argon after cooling to room temperature. To the flask were added 2-[5-(adamantan-1-yl)-*1H*-indol-3-yl]acetic acid (12.4 mg, 0.04 mmol), 2,5-dimethoxy-2'-nitrobenzhydrol (11.6 mg, 0.04 mmol), DMAP (6.1 mg, 0.05 mmol), dry $CH_2Cl_2$ (0.5 mL), and EDC (8.8 μL, 0.05 mmol) at room temperature. The reaction mixture was stirred at room temperature for 16 h. The mixture was quenched with water, and was extracted with EtOAc (3 times). The combined extracts were dried over $Na_2SO_4$, filtered, and evaporated under reduced pressure. The residue was purified by silica-gel column chromatography (hexane : EtOAc = 10 : 1 to 3 : 1) to give caged 5-Ad-IAA (12.5 mg, 54%) as a pale yellow oil. $^1$H NMR (600 MHz, $CDCl_3$) δ 1.72 (d, $J$ = 11.4 Hz, 3H), 1.77 (d, $J$ = 11.4 Hz, 3H), 1.89 (s, 6H), 2.06 (s,

3H), 3.34 (s, 3H), 3.66 (s, 3H), 3.85 (d, $J$ = 15.6 Hz, 1H), 3.89 (d, $J$ = 15.6 Hz, 1H), 6.41 (d, $J$ = 3.0 Hz, 1H), 6.72-6.76 (m, 2H), 7.12 (d, $J$ = 3.0 Hz, 1H), 7.24-7.29 (m, 3H), 7.35-7.40 (m, 2H), 7.54 (s, 1H), 7.73 (s, 1H), 7.91 (dd, $J$ = 7.8, 3.0 Hz, 1H), 8.00 (s, 1H). $^{13}$C NMR (150 MHz, $CDCl_3$) δ 29.0, 31.4, 36.0, 36.8, 43.7, 55.2, 56.0, 68.0, 108.1, 110.6, 111.8, 112.8, 113.8, 114.4, 120.0, 123.3, 124.5, 127.0, 127.9, 128.5, 129.3, 132.7, 134.3, 143.1, 148.4, 150.9, 153.4, 170.2 (1 aryl carbon signal is obscured). HRMS (ESI) $m/z$ calcd for $C_{35}H_{36}O_6N_2Na$ $[M+Na]^+$: 603.2466, found 603.2463.

$^1$H-NMR (600 MHz, $CDCl_3$) of caged 5-Ad-IAA

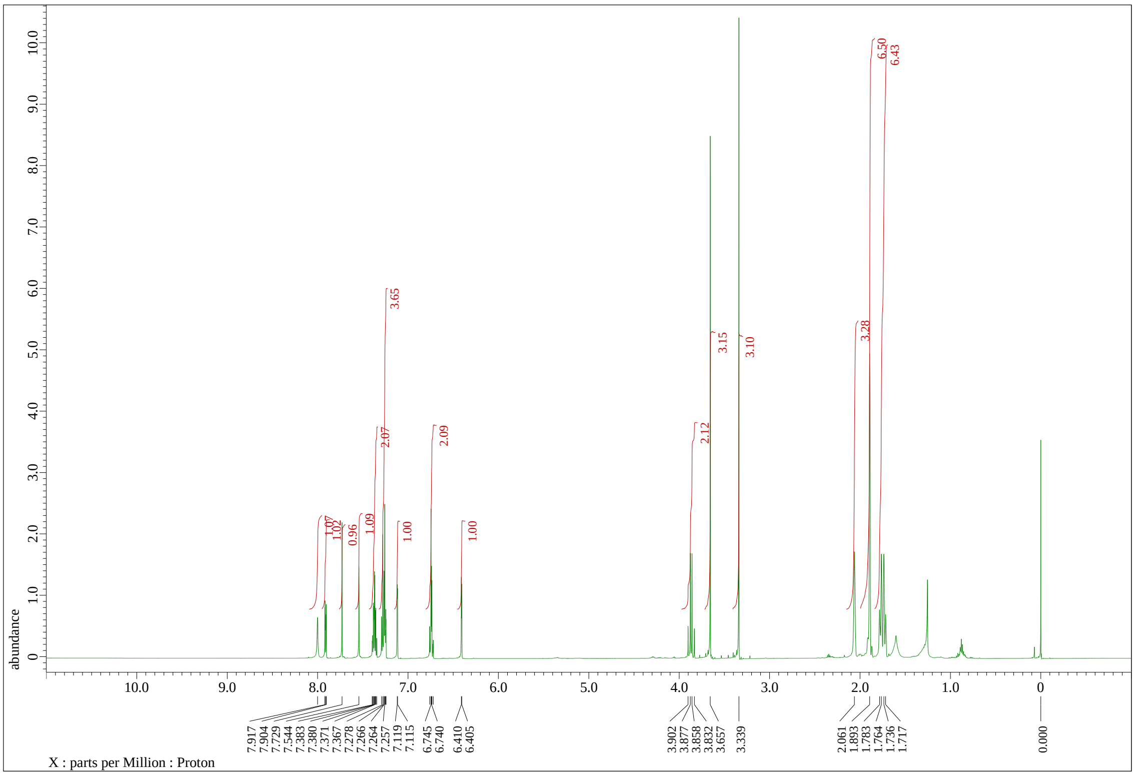

$^{13}$C-NMR (150 MHz, $CDCl_3$) of caged 5-Ad-IAA

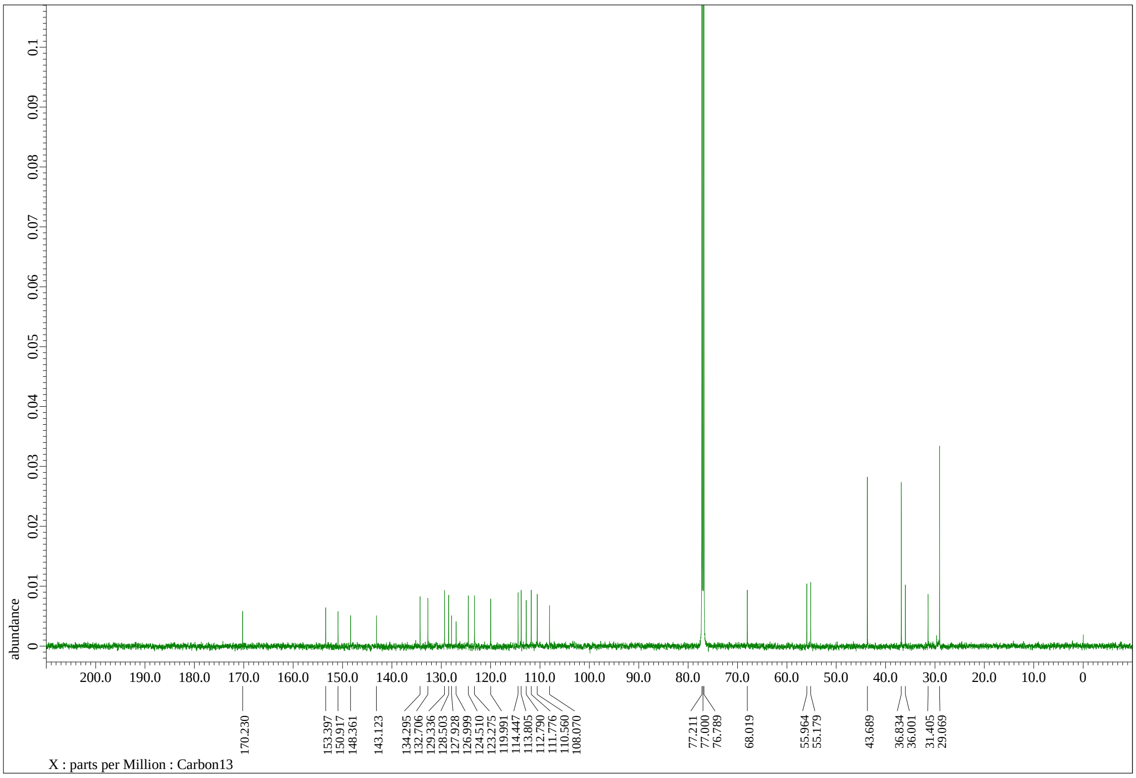
